# Description of a new genus of the *Pectobacteriaceae* family isolated from lake water in France; *Prodigiosinella aquatilis* gen. nov. sp. nov. includes two subspecies *Prodigiosinella aquatilis* subsp. *aquatilis* ssp. nov. and *Prodigiosinella aquatilis* subsp. *natabilis* ssp. nov

**DOI:** 10.1101/2023.09.17.558112

**Authors:** Nicole Hugouvieux-Cotte-Pattat, Jean-Pierre Flandrois, Jérôme Briolay, Sylvie Reverchon, Céline Brochier-Armanet

## Abstract

The *Pectobacteriaceae* family comprises plant pathogens able to provoke diverse diseases, including plant maceration due to the production of pectinases disrupting the plant cell wall. To better understand their natural diversity, a survey of pectinolytic bacteria was performed in lakes of the French region La Camargue near the Mediterranean Sea. Sixteen atypical pectinolytic isolates were obtained from brackish water of three lakes. The genome of six isolates was sequenced; their size is around 4.8 to 5.0 Mb, including a plasmid of 59 to 61 kb; their G+C values range from 49.1 to 49.3 mol%. Phylogenetic analyses indicated that the novel strains formed a new clade of *Pectobacteriaceae*, separate from previously described genera of this family. These analyses suggested also that *Acerihabitans* does not belong to *Pectobacteriaceae* and should be reclassified in the *Bruguierivoracaceae* family, while *Symbiopectobacterium* could be a true *Pectobacteriaceae* member. Based on phenotypic, genomic and phylogenetic characteristics, we propose the creation of a new genus with the name *Prodigiosinella* gen. nov. Both the phenotypic and phylogenetic analyses separated the strains into two distinct subgroups. However, the DNA–DNA relatedness values revealed a close relationship between the two groups, supporting their appurtenance to the same species. Thus, it is proposed to classify them as two subspecies of *Prodigiosinella aquatilis* sp. nov., for which we propose the name *Prodigiosinella aquatilis* subsp. *aquatilis* ssp. nov. (LS101^T^ = CFBP 8826^T^ = LMG 32072^T^) and *Prodigiosinella aquatilis* subsp. *natabilis* ssp. nov. (CE70^T^ = CFBP 9054^T^ = LMG 32867^T^).

## INTRODUCTION

The bacterial order *Enterobacterales* has been created in 2016 [1] and is currently divided into eight recognized families: *Budviciaceae*, *Enterobacteriaceae*, *Erwiniaceae*, *Hafniaceae*, *Morganellaceae*, *Pectobacteriaceae*, *Thorselliaceae*, and *Yersiniaceae*, and the not yet recognized *Bruguierivoracaceae* [1, 2, 3]. Among them, *Pectobacteriaceae* currently comprises the eight genera *Acerihabitans* [4], *Affinibrenneria* [5], *Brenneria* [6], *Dickeya* [7], *Lonsdalea* [6], *Musicola* [8], *Pectobacterium* [9], and *Samsonia* [10], some of them being major plant pathogens. In addition, the genus *Symbiopectobacterium* was recently described as being closely related to *Pectobacterium* [11]. While members of *Brenneria* and *Lonsdalea* are responsible for tree diseases [6], *Dickeya, Musicola*, and *Pectobacterium* affect a wide range of hosts, provoking soft-rot diseases due to the action of extracellular pectinases that degrade the plant cell wall [12, 13]. Most of the characterized strains of these three soft-rot genera were isolated from diseased crops or ornamental plants, but some strains have also been isolated from surface water [14, 15, 16].

In this study, we describe sixteen new bacterial strains isolated from brackish water of lakes in the French region of La Camargue, near the Mediterranean Sea. These strains were selected on the basis of their pectinolytic activity using a semi-selective solid medium [17]. Based on phenotypic and phylogenetic arguments, these isolates were shown to belong to a new genus within *Pectobacteriaceae*, for which we propose the name *Prodigiosinella* gen. nov. These strains can be divided into two phenotypic groups with on one side, LS101^T^, LS102, LS111, LS112, CE90, C110, C111, and on the other side, CE70^T^, C52, C73, C76, C77, C80, C81, C83, C84. The genome sequences of six *Prodigiosinella* strains were determined and phylogenetic studies were performed to refine their classification inside the *Pectobacteriaceae*. Sequence database surveys disclosed other *Prodigiosinella* members isolated in Korea and USA, including the strain ATCC 39006 recently proposed as ‘Candidatus *Prodigiosinella confusarubida*’ [18]. Considering DNA–DNA relatedness values, we propose to classify the *Prodigiosinella* strains in a species named *Prodigiosinella aquatilis* sp. nov., including two subspecies *Prodigiosinella aquatilis* subsp. *aquatilis* ssp. nov. and *Prodigiosinella aquatilis* subsp. *natabilis* ssp. nov., with LS101^T^ (CFBP 8826 ^T^, LMG 32072^T^) and CE70^T^ (CFBP 9054^T^, LMG 32867^T^) as the type strains, respectively.

## STRAIN ISOLATION AND PRELIMINARY CHARACTERIZATION

A survey of pectinolytic bacteria present in lakes of the wetland region La Camargue, near the Mediterranean Sea, in France, revealed atypical isolates corresponding to white or light pink colonies onto crystal violet pectate (CVP) medium [17] (Fig. 1A-B). The water samples were taken from a brackish lake located in a conservation site dedicated to the study and preservation of wetlands (Tour du Valat, 43.5089° N, 4.66757° E, altitude ∼1 m, https://tourduvalat.org/). Our first observation was in June 2020 with four isolates (LS101^T^, LS102, LS111, and LS112) showing, after three days, small deep cavities onto CVP medium, indicating that the colonies were able to degrade the pectate gel [17]. Preliminary analyses based on a few phenotypic tests showed that these four pectinolytic strains constituted an original phenotypic group that clearly differed from members of the pectinolytic genera *Dickeya, Musicola*, or *Pectobacterium* (see below). Furthermore, LS101^T^, LS102, LS111, and LS112 colonies appeared opaque with light pink color on minimal medium and orange-pink color on LB plates, due to the yellow color of LB (Fig. 1B), while *Dickeya, Musicola*, or *Pectobacterium* colonies were translucent and cream (Table 1). A second survey of brackish lakes was performed in October 2020, allowing us to isolate twelve additional strains corresponding to this new phenotypic group (CE90, C110, C111, CE70^T^, C52, C73, C76, C77, C80, C81, C83, C84). The salinity of the corresponding water samples was determined by conductimetry and ranged between 1.93 g.l^-1^ and 6.55 g.l^-1^ in the samples containing the novel type of bacteria.

**Fig. 1.**
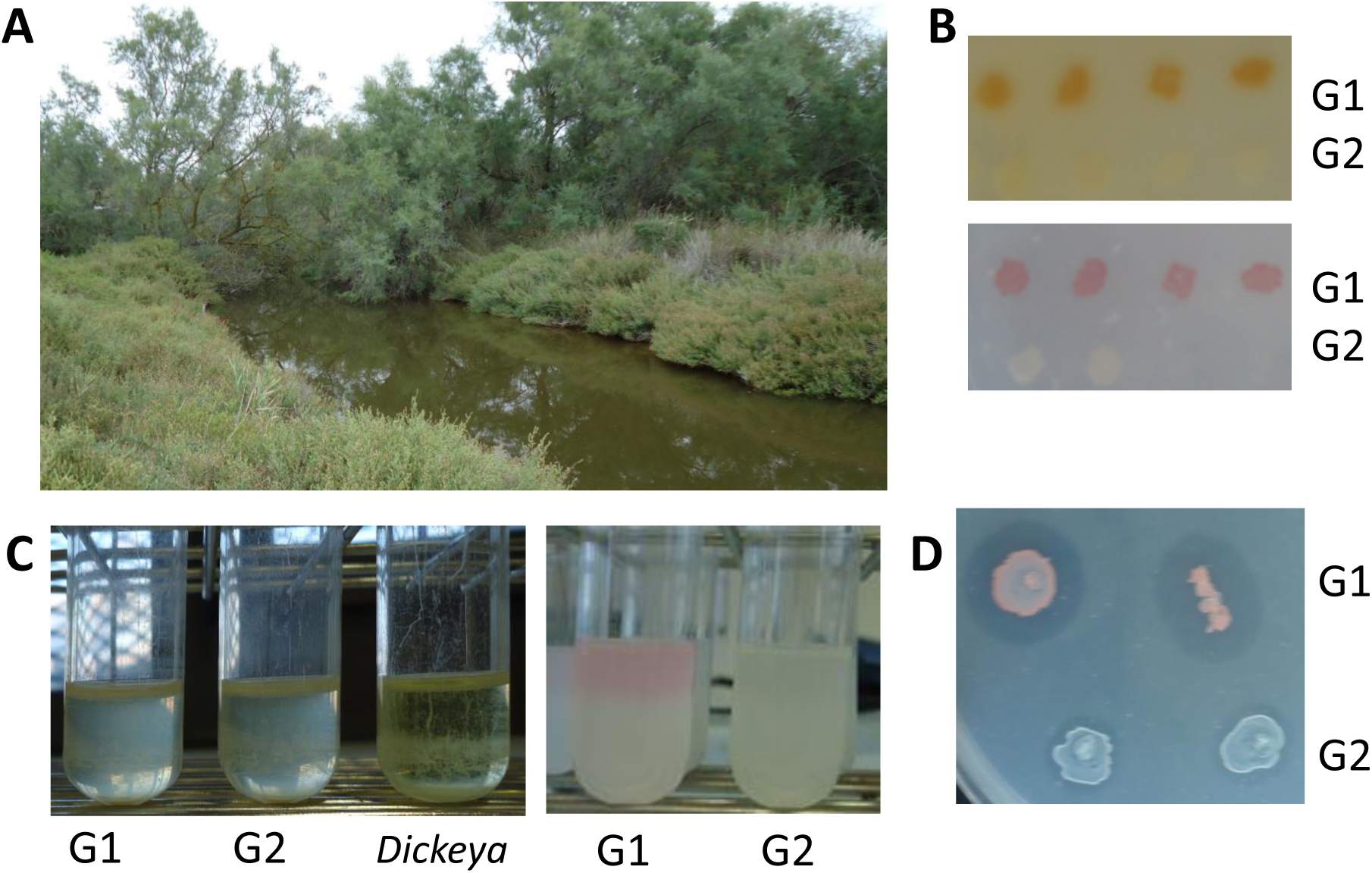
Isolation and characteristics of *Prodigiosinella aquatilis* strains. A, view of the Lake Poutrague (Camargue, France) where the strains were collected. B, color of the colonies on LB and M63 plates. Colonies of the group G1 (*P. aquatilis* subsp. *aquatilis*) appeared opaque with light pink color on minimal medium M63 (bottom) but orange-pink color on LB plates (top), due to the yellow color of the LB medium. Colonies of the group G2 (*P. aquatilis* subsp. *natabilis*) developed a light pink color after at least one or two weeks on M63 medium. C, flotation ability in LB and M63 liquid medium (after growth, cultures were allowed to stand for 7 days; *D. dadantii* 3937 was used as a negative control). D, production of antibacterial compound(s) on a lawn of *E. coli* BW25113 (Δ*tolC*::Kan).

**Table 1.** Phenotypic analysis of the *Prodigiosinella* isolates. The salinity of the water samples was determined by conductimetry (ND = not done). Sugar assimilation was analyzed using strains belonging to the two *Prodigiosinella* groups G1 and G2; a few SRP and *Lonsdalea* strains were used for comparison. Strains were inoculated onto M63 plates supplemented with a sole carbon source (2 g l^-1^) (Dara, D-arabinose; Mel, D-melibiose, Mtl, mannitol; Raf, raffinose; Tre, trehalose; Xyl, D-xylose; PGA, polygalacturonate). Bacterial growth was observed after 24 to 72 hours at 30°C. The sign +, indicates growth; - indicates no growth after 72h at 30°C.

| Strains | Collection names | Classification | Sample salinity | Sugar assimilation |  |  |  |  |  |  | Colony colour |
| --- | --- | --- | --- | --- | --- | --- | --- | --- | --- | --- | --- |
|  |  |  | g.l<sup>-1</sup> | Mtl | D-ara | Xyl | Tre | Mel | Raf | PGA |  |
| LS101<sup>†</sup> | A6420, CFBP8826, LMG32072 | *Prodigiosinella* G1 | ND | + | + | + | + | - | - | + | opaque light pink |
| LS111 | A6421 | *Prodigiosinella* G1 | ND | + | + | + | + | - | - | + | opaque bright pink |
| C111 | A6483, CFBP9052 | *Prodigiosinella* G1 | 6.31 | + | + | + | + | - | - | + | opaque pink |
| CE90 | A6482, CFBP9051 | *Prodigiosinella* G1 | 6.55 | + | + | + | + | - | - | + | opaque bright pink |
| C52 | A6477, CFBP9053 | *Prodigiosinella* G2 | 1.93 | + | + | + | + | + | + | + | opaque white |
| CE70<sup>†</sup> | A6479, CFBP9054, LMG32867 | *Prodigiosinella* G2 | 4.38 | + | + | + | + | + | + | + | opaque light pink |
| C77 | A6478 | *Prodigiosinella* G2 | 4.38 | + | + | + | + | + | + | + | opaque white |
| C80 | A6480, CFBP9055 | *Prodigiosinella* G2 | 3.62 | + | + | + | + | + | + | + | opaque light pink |
| C84 | A6481 | *Prodigiosinella* G2 | 3.62 | + | + | + | + | + | + | + | opaque light pink |
| 3937 | A4922, CFBP3855, NCPPB4426 | *Dickeya dadantii* |  | + | + | + | - | + | + | + | translucent cream |
| S29<sup>†</sup> | A6067, CFBP8647, LMG30899 | *Dickeya lacustris* |  | + | - | - | - | - | - | + | translucent cream |
| NCPPB2511<sup>†</sup> | A6065, CFBP4178 | *Musicola paradisiaca* |  | - | + | + | + | + | + | + | translucent pale cream |
| A3967<sup>†</sup> | CFBP8732<sup>†</sup>, LMG31880<sup>†</sup> | *Musicola keenii* |  | - | + | + | + | - | - | + | translucent pale cream |
| CFBP 8637<sup>†</sup> | A6647<sup>†</sup> | *Pectobacterium aquaticum* |  | + | - | + | + | + | + | + | translucent cream |
| CFBP 6051<sup>†</sup> | A6644<sup>†</sup> | *Pectobacterium versatile* |  | + | - | + | + | + | + | + | translucent cream |
| A6242<sup>†</sup> | CFBP3617<sup>†</sup> | *Lonsdalea quercinia* |  | + | - | - | + | - | - | + | translucent pale cream |

The *gapA* gene is commonly used for preliminary identification in *Pectobacteriaceae* [19]. PCR amplifications were performed on bacterial cell lysates of the sixteen selected isolates using an Illustra PuReTaq Ready-To-Go kit (GE Healthcare) with primers gapA-7-F and gapA-938-R [19]. The *gapA* amplicon sequences were determined by Sanger sequencing (Microsynth France). A BLASTN search against the nr/nt NCBI database gave the *gapA* sequence of strain *Serratia* sp. ATCC 39006 (GCF_000463345.2) as best hit. Recent phylogenomic analyses showed that this strain robustly groups with the genus *Dickeya* within *Pectobacteriaceae* and not with the *Serratia* genus within *Yersiniaceae* [18]. A *gapA* maximum likelihood (ML) tree was constructed for a preliminary classification of the sixteen isolates; it includes 97 strains: the sixteen Camargue isolates, strain ATCC 39006, one strain of each currently recognized *Pectobacteriaceae* species and 18 additional strains from other *Enterobacterales* families. A codon alignment has been built using SEAVIEW 5.0.5 [20] and MUSCLE v3.8.1551 [21] (996 nucleotide positions) and used to infer a ML tree using IQ-TREE v.2.1.2 [22], with the best suited model (SYM+R6) according to ModelFinder [23].

The resulting *gapA* tree shows that the 16 Camargue strains form a monophyletic group (Bootstrap value (BV) = 100%), well separated from other *Pectobacteriaceae* lineages (Fig. 2). The 16 isolates can be further subdivided into two closely related groups, referred hereafter as to G1 and G2 (Fig. 2). G1 contains seven strains (LS101^T^, LS102, LS111, LS112, CE90, C110, and C111) (BV = 100%), while G2 encompasses the nine other strains (C52, C73, C76, C77, CE70^T^, C80, C81, C83, C84) (BV = 94%) (Fig. 2). At the phenotypic level, G1 and G2 strains differ in their ability to use D-melibiose and D-raffinose as carbon sources for growth (Table 1).

**Fig. 2.**
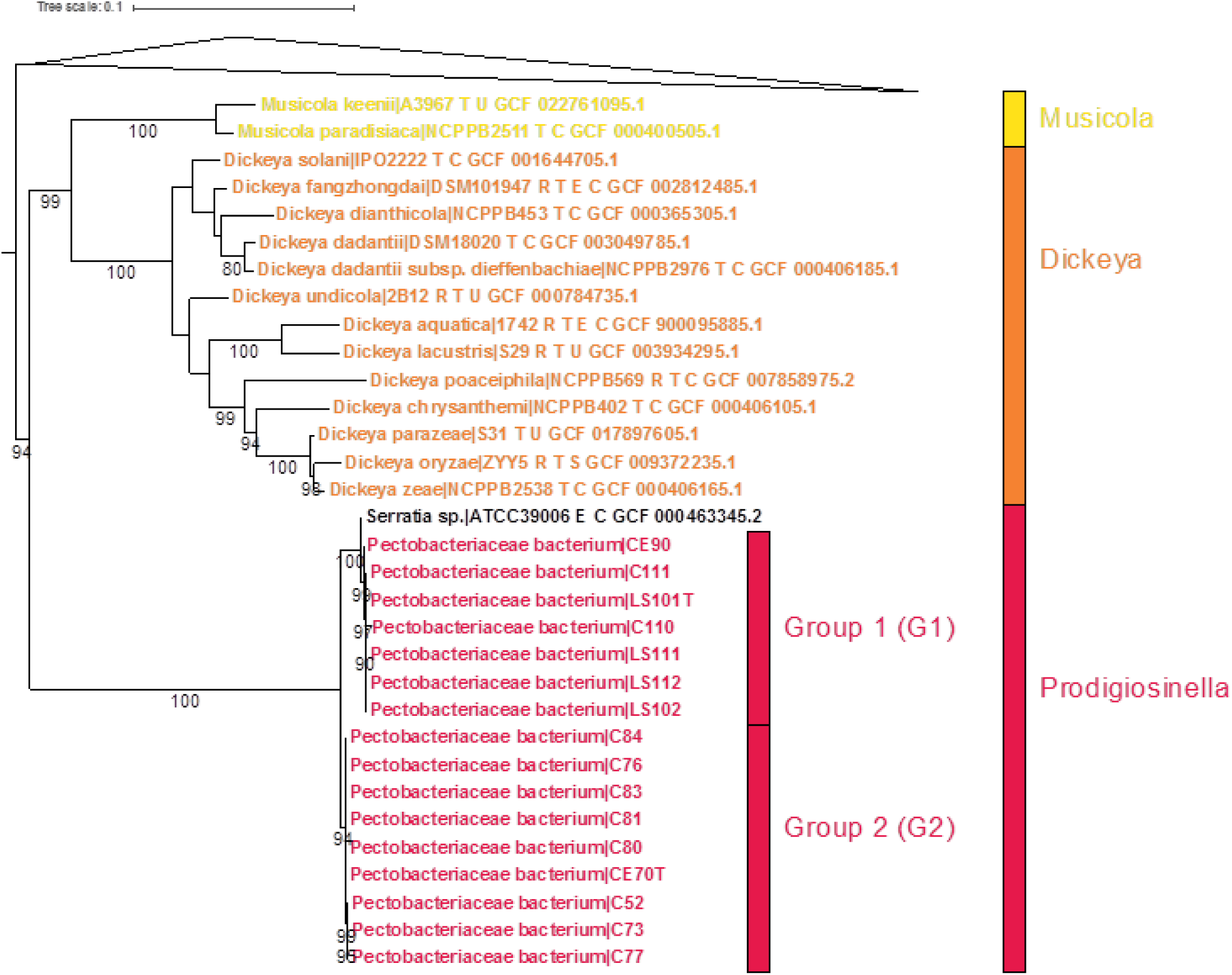
Maximum likelihood phylogeny of the *gapA* gene showing the relationships between the 16 isolates, strain ATCC 39006, and their closest relatives within *Pectobacteriaceae*. The tree has been inferred with IQ-TREE (SYM+R6 model). For clarity, the part of the tree encompassing the sequences of the 16 isolates (in red), strain ATCC 39006 (in black), and their closest relatives (*Dickeya* in orange and *Musicola* in yellow) is detailed, while other *Pectobacteriaceae* sequences have been collapsed (grey triangle). Numbers at branch correspond to ultrafast bootstrap values computed by IQ-TREE (1,000 replicates). For clarity, values <80% are not shown. The scale bar represents the average number of substitutions per nucleotide site.

The 16S rRNA genes of each of the 16 strains were amplified by PCR using primers 27f and 1492r [24] and sequenced (Microsynth France). A BLASTN search against the nr/nt NCBI database showed best-hits with sequences from four strains classified as unidentified species of the genera *Serratia* (ATCC 39006), *Dickeya* (K61), and *Erwinia* (MK01 and MK09). The strain *Serratia* sp. ATCC 39006 was isolated in 1982 from channel water in Cheesequake Salt Marsh (New Jersey, USA), after selection for the production of antibiotics of the carbapenem family [25] and its genome was sequenced in 2013 (GCF_000463345.2) [26]. A new designation was recently proposed for this strain, ‘Candidatus *Prodigiosinella confusarubida*’ [18]. Strains *Erwinia* sp. MK01 and MK09 were isolated in 2004 from the rhizosphere of reeds growing in salt marshes in Korea (AY690711 and AY690717, respectively), while strain *Dickeya* sp. K61 was isolated from freshwater mud in Hawaii (USA) in 2018 (MH894269). The 16S rRNA gene sequences of the 16 Camargue isolates, ATCC 39006, K61, MK01, MK09, and a representative set of *Pectobacteriaceae* type strains were aligned using MAFFT v7.453 with the accurate option L-INSI [27], trimmed with BMGE v.2.0 [28], and used to infer a ML tree using IQ-TREE v2.1.2 [22] with the best suited model (TN+F+I+G4 model) according to ModelFinder [23]. The resulting tree strongly supports the grouping of the Camargue isolates with strains ATCC 39006, K61, MK01, and MK09 (BV = 100%, Fig. 3). Consistently with the analysis of the *gapA* gene, this group forms a distinct lineage, distant from the other *Pectobacteriaceae* genera, suggesting it represents a novel genus within *Pectobacteriaceae*. As suggested previously [19], we propose the name *Prodigiosinella* gen. nov. for this novel genus, referring to the presence of the *pig* cluster in the genome of ATCC 39006 and in the six sequenced genomes (see below). This gene cluster allows the synthesis of the red pigment prodigiosin, explaining the pink color of the Camargue isolates (Fig. 1B). The geographical repartition of this new genus is large as *Prodigiosinella* members were isolated in four different continents, North America (strain ATCC 39006), Asia (strains MK01 and MK09) Oceania (strain K61) and Europe (this work).

**Fig. 3.**
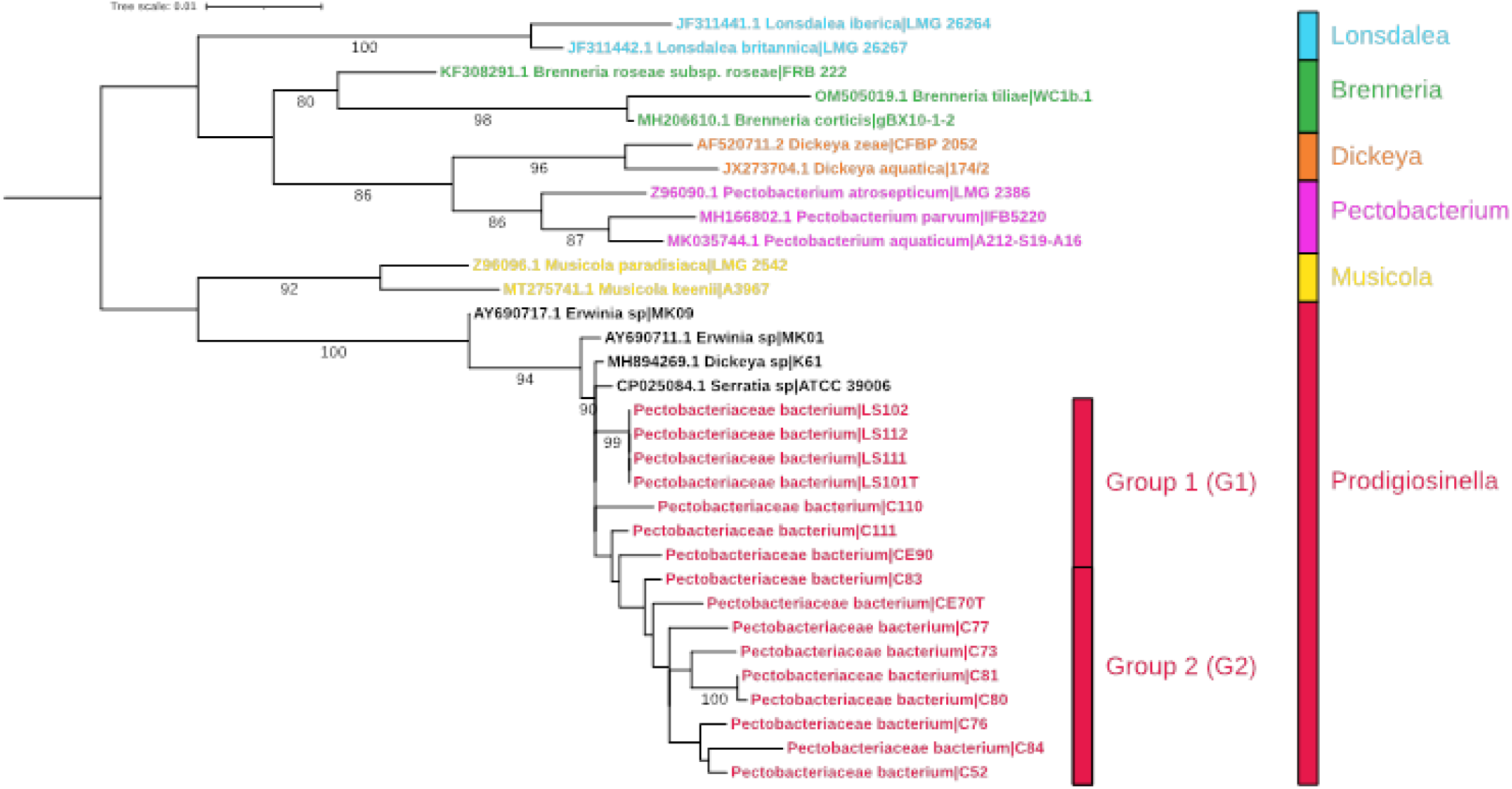
Maximum likelihood phylogeny of *Prodigiosinella* strains based on the 16S rRNA gene sequences. The tree was inferred with IQ-TREE (TN+F+I+G4 model) using sequences of the 16S rRNA PCR products of 16 strains isolated in Camargue and the available 16S rRNA gene sequences of *Serratia* sp. ATCC 39006 (CP025084.1), *Erwinia* sp. MK01 and MK09 (AY690711 and AY690717), *Dickeya* sp. K61 (MH894269), and a few *Pectobacteriaceae* type strains (32 sequences, 1,468 nucleotide sites). The scale bar represents the average number of substitutions per site. Numbers at branch represent ultrafast bootstrap values computed with IQ-TREE (1,000 replicates). For clarity values <80% are not shown.

## PHENOTYPIC ANALYSIS

A biochemical characterization was performed using four *Prodigiosinella* strains from G1 (LS101^T^, LS111, C111, CE90) and five from G2 (C52, C77, CE70^T^, C80, C84) (Table 1). To test their growth with different carbon sources, the strains were inoculated on M63 minimal medium supplemented with the appropriate carbon source. The strains from the groups G1 and G2 differ by their capacity to assimilate melibiose and raffinose (Table 1). In contrast, all strains are able to grow in the presence of D-arabinose, D-mannitol, D-trehalose, and D-xylose. These biochemical characteristics are sufficient to distinguish *Prodigiosinella* strains from *Dickeya, Musicola*, and *Pectobacterium* whose members are unable to assimilate D-trehalose, D-mannitol, and D-arabinose, respectively (Table 1) [8, 29, 30]. *Prodigiosinella* strains also differ by several characters from the close genus *Lonsdalea*, such as for the assimilation of D-arabinose and xylose (Table 1).

To complete this phenotypic characterization, we analyzed three *Prodigiosinella* strains of each group (for G1 strains LS101^T^, C111, and CE90; for G2 strains C52, C80, and CE70^T^) using Biolog plates PM1 and PM2A which contain 190 potential carbon sources [31] (Table S1). Inoculations were performed according to the manufacturer instructions and the lecture was made after 48h at 30°C. Rather than bacterial growth, the Biolog system detects the metabolic activity of the cells due to substrate assimilation; this activity is visualized by the color change of an oxidoreduction indicator [31]. The six *Prodigiosinella* strains assimilate the following sugars: N-acetyl-D-glucosamine, D-arabinose, L-arabinose, D-fructose, D-galactose, D-galacturonic acid, D-gluconic acid, D-glucosamine, α-D-glucose, glycerol, m-inositol, D-mannitol, D-mannose, mucic acid, β-methyl-D-glucoside, D-ribose, L-rhamnose, D-saccharic acid, D-sorbitol, D-trehalose, and D-xylose. They assimilate different plant-derived glycosides, oligo or polysaccharides such as arbutin, D-cellobiose, gentiobiose, salicin, sucrose, and pectin. They also metabolize several small organic acids: γ-amino butyric acid, citric acid, fumaric acid, D-malic acid, L-malic acid, malonic acid, pyruvic acid, and succinic acid (Table S1). New differences are observed between G1 and G2 strains. While G1 strains are able to assimilate m-hydroxy phenyl acetic acid, p-hydroxy phenyl acetic acid, tyramine and D.L-octopamine, G2 strains could not. Inversely, only G2 strains are able to use D-melibiose and D-raffinose, and to weakly assimilate D-alanine, L-alanine, thymidine, and D,L-α-glycerol-phosphate (Table S1). These phenotypic differences show that G1 and G2 strains represent two closely but distinct phenotypic groups, corresponding possibly to two *Prodigiosinella* species or subspecies.

The effect of temperature on the growth of six selected *Prodigiosinella* strains was analyzed in LB medium by incubations ranging from 25 to 41°C. Cell density was estimated by measuring optical density at 600 nm (OD_600_) at 24h. Growth of the *Prodigiosinella* strains is observed in a large range, from 25 to 36°C, but not at 39°C (Fig. S1).

The strain ATCC 39006 was originally identified on the basis of its capacity to produce antibiotic [25]. Consistently, the new *Prodigiosinella* isolates are able to inhibit the growth of the *E. coli* strain BW25113 (Δ*tolC*::Kan), more efficiently for G1 strains than for G2 strains (Fig 1D). Since ATCC 39006 was shown to produce gas vesicles allowing bacterial cells to float in stagnant water [32], we tested the flotation ability of the new *Prodigiosinella* isolates. After growth with shaking in liquid medium, cultures were allowed to stand for 7 days at room temperature. While *Dickeya* cells settle at the bottom of the tube, cells of the Camargue isolates remain in suspension throughout the liquid, suggesting that they are able to produce gas vesicles (Fig 1C).

*Dickeya* and *Musicola* are characterized by their ability to secrete several plant cell wall degrading enzymes. Because of their close relationship with *Prodigiosinella*, we investigated these activities in Camargue strains using a range of media [33] (Table 2). In each case, *D. dadantii* 3937 was used as a reference strain. The six *Prodigiosinella* strains show pectinase activity on medium containing polygalacturonate, and they are able to grow in the presence of this polysaccharide as a sole carbon source (Table 1). In contrast, no significant cellulase or protease activity was detected (Table 2). The ability to macerate plant tissues was evaluated by measuring the percentage of inoculations leading to maceration and the quantity of macerated tissue obtained 48 h after inoculation on either chicory leaves or potato tubers [33]. Only a small part of the inoculations with a *Prodigiosinella* strain results in plant tissue maceration and, in these cases, the quantity of macerated tissue remains low (Table 2). Thus, in comparison to *Dickeya* strains, the *Prodigiosinella* strains show a very low efficiency to macerate plant tissues.

**Table 2.** Enzyme secretion, motility and maceration ability. *D. dadantii* 3937 was used as a reference strain. The enzyme secretion was evaluated after 24h growth on plates containing an enzyme substrate allowing the detection of protease (Prt), pectinase (Pel) or cellulase (Cel) activity: +, positive; w, weak; -, negative. Motility was estimated by the growth diameter 24 h after inoculation into 0.3% L agar plate for swimming and on 0.6% L agar plate for swarming. For virulence tests, the percentage of plants presenting maceration was observed 48 h after the bacterial inoculation. The length of macerated tissue (mm) and the weight of macerated tissue (g) were measured to quantify maceration on chicory leaves and potato tubers, respectively. For each measurement, the mean value was calculated only for plants showing maceration.

| Strain | Identification | Secreted enzymes |  |  | Mobility |  | Virulence tests |  |  |  |
| --- | --- | --- | --- | --- | --- | --- | --- | --- | --- | --- |
|  |  | Pel | Cel | Prt | Swimming | Swarming | Chicory leaves |  | Potato tubers |  |
|  |  |  |  |  | mm | mm | % disease | mm | % disease | g |
| **LS101<sup>T</sup>** | *Prodigiosinella* G1 | w | - | - | 35 | 10 | 10 | 25 | 30 | 0.47 |
| **C111** | *Prodigiosinella* G1 | w | - | - | 39 | 12 | 0 | 0 | 20 | 0.15 |
| **CE90** | *Prodigiosinella* G1 | w | - | - | 36 | 30 | 0 | 0 | 10 | 0.28 |
| **C52** | *Prodigiosinella* G2 | w | - | - | 23 | 6 | 20 | 10 | 90 | 0.66 |
| **CE70<sup>T</sup>** | *Prodigiosinella* G2 | w | - | - | 24 | 18 | 20 | 10 | 50 | 0.32 |
| **C80** | *Prodigiosinella* G2 | w | - | - | 19 | 6 | 0 | 0 | 60 | 0.43 |
| **S29<sup>T</sup>** | *D. lacustris* | + | + | + | 28 | 9 | 100 | 32 | 100 | 1.55 |
| **3937** | *D. dadantii* | + | + | + | 31 | 85 | 100 | 89 | 100 | 4.67 |

The bacterial motility was measured by the growth diameter 24h after inoculation in 0.3% L agar plate for swimming and on 0.6% L agar plate for swarming [33]. In comparison to *D. dadantii*, the six *Prodigiosinella* strains show a high swimming motility but a moderate swarming activity (Table 2).

## GENOME SEQUENCES AND GENOMIC RELATEDNESS

The genome of six *Prodigiosinella* strains was sequenced (for G1: LS101^T^, C111, CE90; for G2: C52, C80, CE70^T^). DNA extraction was performed using a New England Biolabs kit (https://international.neb.com/monarch/high-molecular-weight-dna-extraction). Genomic DNA was sequenced by combining both the Illumina technique (Microbes NG, UK) and a Nanopore Minion apparatus in the platform DTAMB (Federation BioEEnViS, FR3728 UCBL-CNRS). Nanopore reads were filtered by using FiltLong (https://github.com/rrwick/Filtlong), removing the 20% low quality reads and reads of length less than 5,000 bp. Retained reads were then assembled by using Flye v2.9.1 [34]. The circularity of the assembly was verified with Bandage v0.9.0 [35]. The Illumina reads were aligned on the assembly with Bowtie 2 v2.5.0 [36], and indexed and sorted with Samtools v1.17 [37], and used to polish the Nanopore assembly with Pilon v1.23 [38].

Complete genome sequences were obtained for the six strains (Table 3). The six strains contain a circular plasmid. The plasmid of G1 strains, pPrAq1, is slightly smaller (∼59 kb) than pPrAq2 found in G2 strains (∼61 kb). In contrast, no plasmid was reported in the ATCC 39006 genome (Table 3). The size of *Prodigiosinella* genomes (including ATCC 39006) is around 4.8-5 Mb, in the same range as *Dickeya* or *Musicola* genomes (4.3-5.35 and 4.4-4.68 Mb, respectively) but higher than the *Lonsdalea* genomes ranging from 3.54 to 4.02 Mb. The *Prodigiosinella* genomes have G+C contents of 49.1 to 49.3 mol%, clearly lower than that of *Dickeya*, *Musicola* or *Lonsdalea* genomes (52.6 to 56.9 mol%). The taxonomic relations between the *Prodigiosinella* strains were investigated by calculation of *in silico* DNA–DNA hybridization (dDDH) and average nucleotide identity (ANI) values [39, 40] (Table 4). dDDH and ANI values were calculated based on pairwise comparisons between the genomes of the six Camargue strains and strain ATCC 39006 (Table 4). dDDH and ANI values among *Prodigiosinella* strains are high (>73% for dDDH and >97% for ANI) (Table 4). Thus, based on a dDDH threshold of 70% and an ANI threshold of 95% to separate two species [39, 40], all the strains of the proposed new genus *Prodigiosinella* belong to the same species for which we propose the name *Prodigiosinella aquatilis*. However, while dDDH values are >91% within each group of *Prodigiosinella* strain, they are < 77% between the two groups G1 and G2. Considering a threshold dDDH value of 79-80% for delineating subspecies [41], each group may correspond to a subspecies for which we propose the names *Prodigiosinella aquatilis* subsp. *aquatilis* ssp. nov. and *Prodigiosinella aquatilis* subsp. *Natabilis* ssp. nov.

**Table 3.** Characteristics and accession numbers of the *Prodigiosinella* genomes.

|  | *Prodigiosinella aquatilis aquatilis* |  |  | *Prodigiosinella aquatilis natabilis* 641 |  |  |  |
| --- | --- | --- | --- | --- | --- | --- | --- |
| Strains | LS101<sup>†</sup> | C111 | CE90 | C52 | CE70<sup>†</sup> | C80 | ATCC 39006 |
| Genome accession | CP128857 | CP128859 | CP129114 | CP128861 | CP128863 | CP129116 | GCF_000463345.2 |
|  | CP128858 | CP128860 | CP129115 | CP128862 | CP128864 | CP129117 |  |
| BioProject | PRJNA905946 | PRJNA905946 | PRJNA905946 | PRJNA905946 | PRJNA905946 | PRJNA905946 | PRJNA224116 |
| BioSample | SAMN-31888678 | SAMN-31888679 | SAMN-31888680 | SAMN-31888681 | SAMN-31888682 | SAMN-31888683 | SAMN-02470747 |
| Genome size | 4,918,023 bp | 4,918,635 bp | 4,898,109 bp | 4,941,093 bp | 4,892,228 pb | 4,858,875 pb | 4,971,757 pb |
| Plasmid size | 59,070 bp | 59,071 bp | 59,832 bp | 61,716 pb | 61,717 pb | 61,716 | NA |
| Genomic GC% | 49.1% | 49.1% | 49.2% | 49.1% | 49.1% | 49.1% | 49.3% |
| Genes (total) | 4,644 | 4,638 | 4,574 | 4,638 | 4,518 | 4,509 | 4,519 |
| Genes (CDS) | 4,537 | 4,531 | 4,468 | 4,528 | 4,408 | 4,399 | 4,415 |
| - Coding | 4,357 | 4,362 | 4,270 | 4,306 | 4,291 | 4,195 | 4,310 |
| - Pseudogenes | 180 | 169 | 198 | 222 | 117 | 204 | 105 |
| Genes (RNA) | 107 | 107 | 106 | 110 | 110 | 110 | 104 |
| - rRNAs 5S | 8 | 8 | 8 | 8 | 8 | 8 | 8 |
| - rRNAs 16S | 7 | 7 | 7 | 7 | 7 | 7 | 7 |
| - rRNAs 23S | 7 | 7 | 7 | 7 | 7 | 7 | 7 |
| - tRNAs | 75 | 75 | 75 | 75 | 75 | 75 | 74 |
| - ncRNAs | 10 | 10 | 9 | 13 | 13 | 13 | 8 |
| CRISPR arrays | 1 | 1 | 1 | 1 | 1 | 1 | 4 |

**Table 4.**
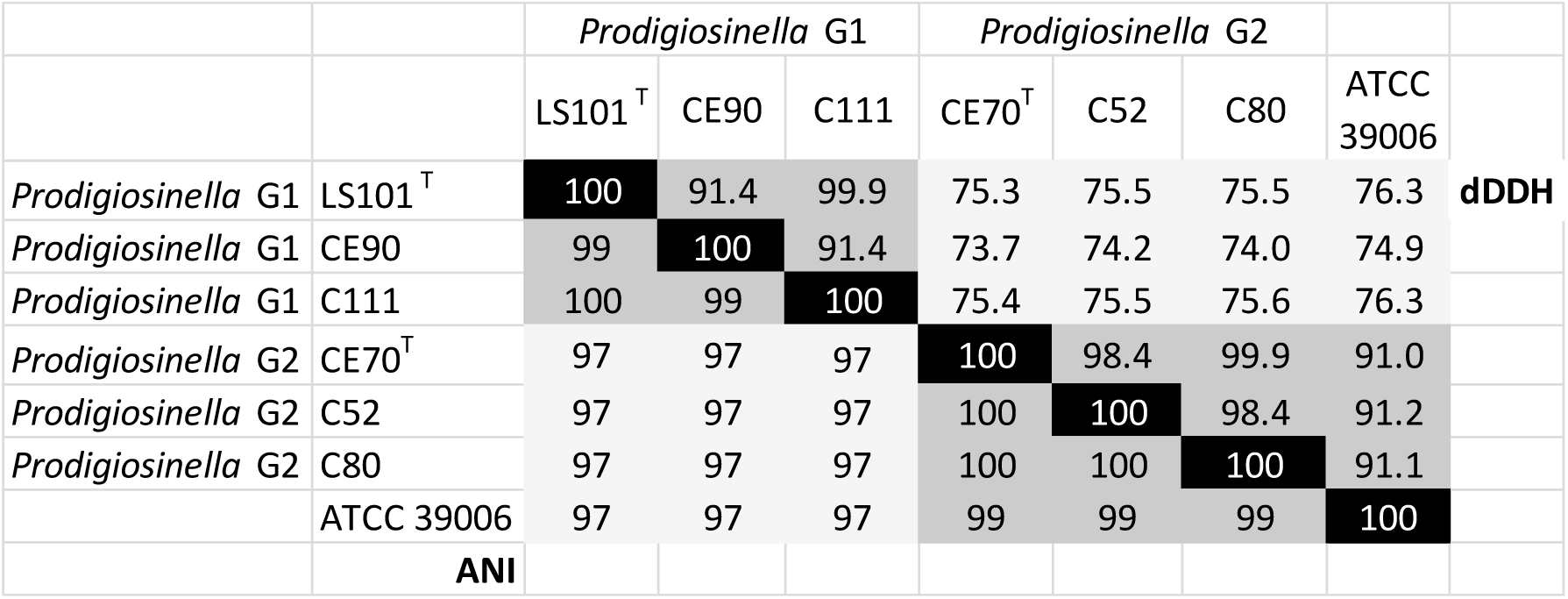
ANI and dDDH values for the genomes of the *Prodigiosinella* isolates. This analysis was performed using the six *Prodigiosinella* genomes of groups G1 or G2 and that of strain ATCC 39006. The ANI values were calculated using the web-service at http://www.ezbiocloud.net/tools/ani. For dDDH calculation, the genomes sequences were uploaded to the Type (Strain) Genome Server (TYGS) (https://tygs.dsmz.de). All pairwise comparisons among the set of genomes were conducted using GBDP and accurate intergenomic distances inferred under the algorithm ‘trimming’ and distance formula *d5*.

|  |  | *Prodigiosinella* G1 |  |  | *Prodigiosinella* G2 |  |  | ATCC 39006 |  |
| --- | --- | --- | --- | --- | --- | --- | --- | --- | --- |
|  |  | LS101<sup>T</sup> | CE90 | C111 | CE70<sup>T</sup> | C52 | C80 |  |  |
| *Prodigiosinella* G1 | LS101<sup>T</sup> | 100 | 91.4 | 99.9 | 75.3 | 75.5 | 75.5 | 76.3 | dDDH |
| *Prodigiosinella* G1 | CE90 | 99 | 100 | 91.4 | 73.7 | 74.2 | 74.0 | 74.9 |  |
| *Prodigiosinella* G1 | C111 | 100 | 99 | 100 | 75.4 | 75.5 | 75.6 | 76.3 |  |
| *Prodigiosinella* G2 | CE70<sup>T</sup> | 97 | 97 | 97 | 100 | 98.4 | 99.9 | 91.0 |  |
| *Prodigiosinella* G2 | C52 | 97 | 97 | 97 | 100 | 100 | 98.4 | 91.2 |  |
| *Prodigiosinella* G2 | C80 | 97 | 97 | 97 | 100 | 100 | 100 | 91.1 |  |
|  | ATCC 39006 | 97 | 97 | 97 | 99 | 99 | 99 | 100 |  |
|  | ANI |  |  |  |  |  |  |  |  |

## PHYLOGENOMICS SUPPORTING ESTABLISHMENT OF *PRODIGIOSINELLA* GEN. NOV

For these analyses, the six sequenced *Prodigiosinella* strains, strain ATCC 39006, and a set of 50 strains from currently recognized *Pectobacteriaceae* genera (*Acerihabitans*, *Affinibrenneria*, *Brenneria*, *Lonsdalea*, *Dickeya*, *Musicola*, *Pectobacterium*, *Symbiopectobacterium*, and *Samsonia*) were considered. The outgroup was composed of 30 representatives of other *Enterobacterales* families. The complete genomes of the 87 corresponding strains were retrieved from NCBI. Genomic sequences were annotated using the NCBI Prokaryotic Genome Annotation Pipeline (PGAP) v.2022-12-13.build6494 [42]. Gene families were delineated according to information from the annots.gff files generated by PGAP (i.e. gene name, function, and hmm profile number when available). The core protein families are defined as the set of 737 protein families with exactly one homologue in 95% of the 87 considered genomes. Ribosomal proteins (rprots) were retrieved from the RiboDB database [43]. The 51 rprot families present in more of 80% of the 87 genomes were kept.

Amino acid sequences of core protein families were aligned using MAFFT v7.453 with the accurate option L-INSI [27], while nucleotide sequences of rprots genes were aligned using SEAVIEW 5.0.5 [20] and MUSCLE v3.8.1551 [21]. The resulting multiple alignments were trimmed using BMGE v.2.0 [28]. Rprots gene alignments on the one hand and core protein alignments on the other hand were then combined to build two large supermatrices containing, respectively, 19,311 nucleotide and 221,882 amino acid positions. The ML trees of ribosomal genes and core proteins were inferred with the SYM+I+I+R5 and the Q.insect+F+I+I+R9 evolutionary models, respectively, as suggested by ModelFinder. Branch supports were estimated with ultrafast bootstrap (1,000 replicates) for the ribosomal gene supermatrix and a jackknife sampling of 7.5% of the core protein supermatrix sites (1,000 replicates). ML trees inferred with these two supermatrices support the monophyly of *Pectobacteriaceae*, with the exception of *Acerihabitans* (Fig. 4 and Fig. S2). In fact, this genus robustly branches with *Sodalis* and *Biostraticola* (Jackniffe value (JV) and BV = 100%). This is consistent with the original publication describing *Acerihabitans* as a new genus of *Pectobacteriaceae*, close to *Sodalis* and *Biostraticola* [4]. The same year, Li and colleagues suggested that *Sodalis* and *Biostraticola* are not *Pectobacteriaceae*, but the members of a new family of *Enterobacterales*, tentatively called *Bruguierivoracaceae* [3]. However, *Acerihabitans* representatives were not included in their analysis, which likely explains why this genus is still considered as part of *Pectobacteriaceae*. Our results indicate clearly that *Acerihabitans* is close to *Sodalis* and *Biostraticola* and consistently, should be reclassified as a member of *Bruguierivoracaceae*. The ML trees of ribosomal genes and core proteins also strongly support the relationship between the *Symbiopectobacterium* genus and *Pectobacteriaceae* (JV and BV = 100%, Fig. 4 and Fig. S2), suggesting that *Symbiopectobacterium* is likely part of this family.

**Fig. 4.**
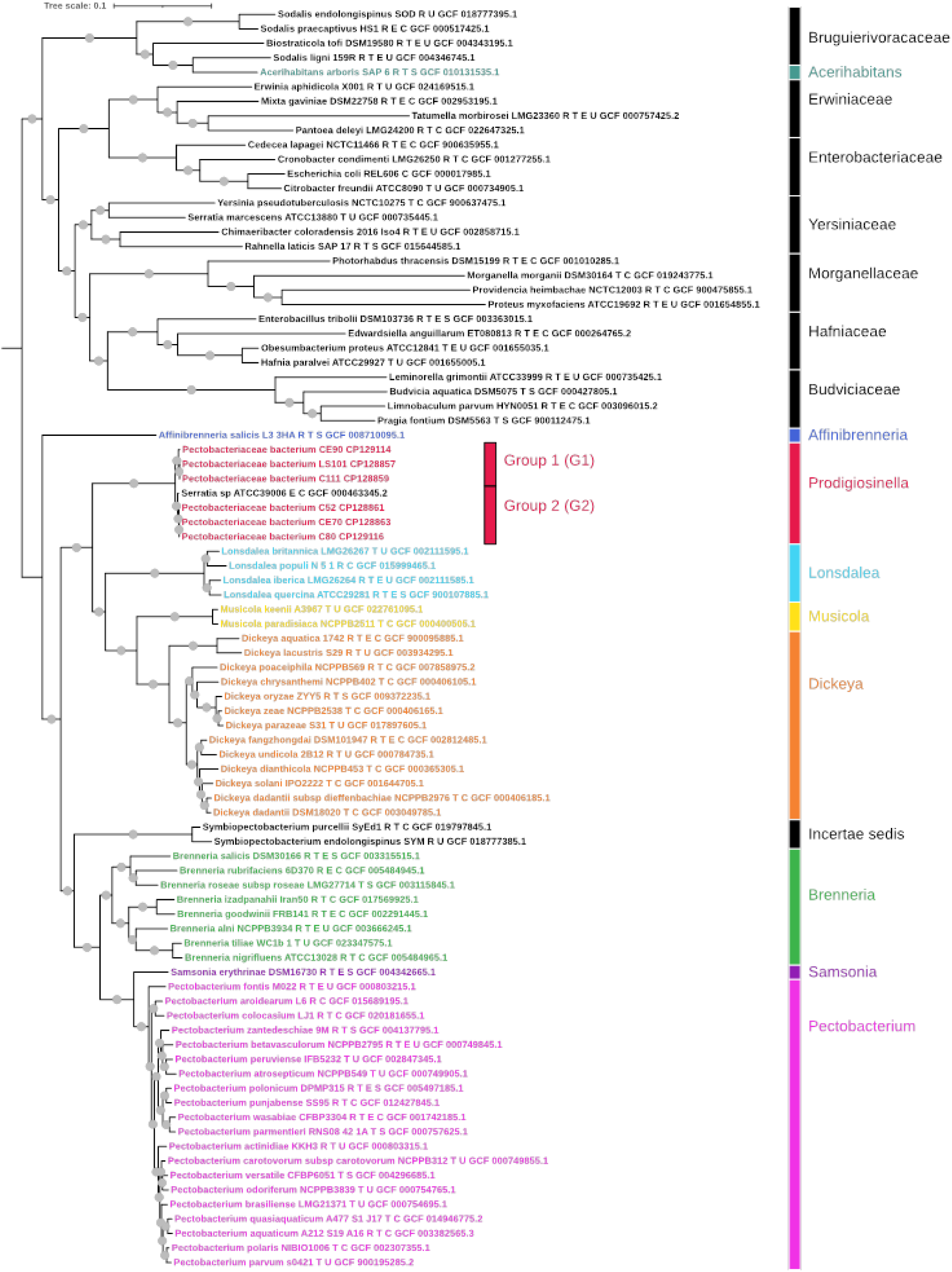
Maximum likelihood phylogeny of the *Pectobacteriaceae* based on core proteins. The tree shows the relationships between the six Camargue isolates, strain ATCC 39006, and *Pectobacteriaceae* genera (87 genome sequences, 221,882 amino acid positions). The tree is rooted with a sample of other *Enterobacterales* families. The tree was inferred with IQ-TREE (Q.insect+F+I+I+R7 evolutionary model, as suggested by IQ-TREE) through a jackknife procedure (1,000 replicates, random selection of 7.5% of the positions). The scale bar represents the average number of substitutions per site. Grey nodes at branch correspond to jackknife supports = 100%.

Consistently with the *gapA* tree, the six *Prodigiosinella* isolates form a well-supported monophyletic group that also includes ATCC 39006 (JV and BV = 100%, Fig. 4 and Fig. S2). This group is clearly separated from other *Pectobacteriaceae* genera and branches as the sister-lineage of the large clade encompassing *Lonsdalea*, *Musicola*, and *Dickeya*, three major phytopathogenic genera (BV = 78% and JV = 100%). Regarding the relationships between the strains, both trees strongly recover the G1 and G2 dichotomy between strains, with ATCC 39006 belonging to the G2 group (BV and JV = 100%). Altogether, the phylogenetic analyses strongly sustain that Camargue isolates and ATCC 39006 represent a new genus within *Pectobacteriaceae*.

## FUNCTIONAL GENOMICS

Genome comparisons were used to identify distinctive traits of the *P. aquatilis* strains. First, we looked for the presence of two gene clusters encoding the biosynthesis of secondary metabolites previously described in ATCC 39006. The *pig* cluster (*pigABCDEFGHIJKLMN*) allows the biosynthesis of the red pigment prodigiosin (2-methyl3-pentyl-6-methoxyprodigiosin) [44], and the *car* cluster (*carABCDEFGH*) codes for genes involved in the synthesis of the carbapenem antibiotic 1-carbapen-2-em-3-carboxylic acid [45] (Table S2). The six sequenced genomes contain the two complete clusters, explaining the pink color of the colonies and the inhibition of *E. coli* growth, respectively (Fig. 1B and 1D). Among *Enterobacterales*, the *pig* cluster and prodigiosin production were observed only in a few *Serratia* species [47]. In contrast, the *car* cluster is quite widespread and found in some strains of several *Pectobacteriaceae* genera, such as *Brenneria*, *Dickeya*, *Musicola*, *Pectobacterium*, *Samsonia*, and *Symbiopectobacterium*. The seven *P. aquatilis* genomes also contain the *gvp* gene cluster (*gvpA_1_CNVF_1_GWA_2_KXA_3_Y-gvrA-gvpHZF_2_F_3_gvrBC*) involved in the production of gas vehicles [46] responsible for the capacity of cell flotation in liquid medium and the opaque aspect of the colonies (Fig. 1). Cell flotation is a rare property and the *gvp* cluster is not found in any other *Enterobacteral*es genome.

Some differences between the genomes of the two *P. aquatilis* subspecies are observed, in particular the presence of genes *rafAT* involved in both melibiose and raffinose assimilation in the genomes of *P. aquatilis* subsp*. natabilis*, including ATCC 39006 (group G2). This explains the phenotypic differences observed between the G1 and G2 strains for melibiose and raffinose assimilation (Table 1). The differences regarding the assimilation of hydroxyphenyl acetic acids can be explained by the presence in *P. aquatilis* subsp*. aquatilis* (group G1) of the *hpa* gene cluster (*hpaRGEDFJSHIXABC*) involved in the catabolism of these compounds [48, 49](Table S1, Table S2). Compared to the cluster *hpa* of *E. coli* strains degrading hydroxyphenyl acetic acids [48], the *P. aquatilis* cluster encodes the additional protein HpaJ. This cluster is also involved in the catabolism of tyramine and D.L-octopamine [49], two substrates which are assimilated by *P. aquatilis* subsp*. aquatilis* (Table S1).

A survey of *P. aquatilis* gene annotations was performed to search for genes potentially involved in the degradation of plant cell walls, and especially pectate lyases, the main determinants of the soft rot symptoms caused by soft-rot *Pectobacteriaceae* [50] (Table S2, Table 5). The seven *P. aquatilis* genomes contain two genes encoding pectate lyases of the family PL1 (PelB, PelD), one of family PL9 (PelX), an intracellular oligogalacturonate lyase of the family PL22 (Ogl), an hydrolase of the family GH105 (RhiN), a pectin methylesterase of the family CE8 (PemA), two pectin acetylesterases of family CE12 (PaeX and PaeY), an endogalactanase of the family GH53 (GanA) and an exogalactanase of the family GH42 (GanB) (PL, GH and CE families are defined on CAZy (http://www.cazy.org/) [51]. All the genomes, except that of C80, encode a pectate lyase of the family PL2 (PelW). All the genomes, except C52, encode a polygalacturonase of the family GH28 (PehV) and a glucuronoxylanase of the family GH30 (XynA). The four *P. aquatilis* subsp. *natalis* genomes contain additional genes encoding two PL1 pectate lyase (PelC, PelZ), another PL9 pectate lyase (PelN), a PL1 pectin lyase (PnlG), a rhamnogalacturonate lyase of the family PL4 (RhiE), and a periplasmic CE8 pectin methyl esterase (PemB). In *P. aquatilis* subsp. *aquatilis*, there is only a partial sequence of *pelN*, *rhiE* and *pemB* that are annotated as pseudogenes. Thus, *P. aquatilis* subsp. *natalis* genomes contain more genes encoding plant cell walls degrading enzymes that the *P. aquatilis* subsp. *aquatilis* genomes (Table 5). In addition, the *P. aquatilis* subsp. *natalis* strain ATCC 36009 contains two adjacent genes, *pelL-celZ*, encoding a PL9 pectate lyase (PelL) and a cellulase of the family GH5 (CelZ), respectively. Interestingly, such cluster is also present in the *D. dadantii* genome (Table 5).

**Table 5.** Genes or clusters encoding plant cell wall degrading enzymes in *Prodigiosinella aquatilis* Strains. PL, GH and CE enzyme families are defined on CAZy (http://www.cazy.org/). The cellular localization of the corresponding protein in *D. dadantii* is indicated by the colour: red genes encoding secreted enzymes, blue genes encoding periplasmic enzymes, green genes encoding cytoplasmic enzymes, purple genes encoding membrane enzyme, and black genes for unknown protein localization.

| *Dickeya dadantii* 3937<br>genes encoding plant cell<br>wall degrading enzymes | *Prodigiosinella aquatilis*<br>subsp. *aquatilis* strains<br>LS101<sup>T</sup>, CE90, C111 | *Prodigiosinella aquatilis* subsp. *natabilis* |  |
| --- | --- | --- | --- |
|  |  | strains CE70<sup>T</sup>, C52, C80 | strain ATCC 39006 |
| *pelA-pelE-pelD-paeY-pemA*<br>PL1 PL1 PL1 CE12 CE8 | *pelD-paeY-pemA* |  |  |
| *pelX* PL9 | *pelX* |  |  |
| *ogl* PL22 | *ogl* |  |  |
| *rhiN* GH105 | *rhiN* |  |  |
| *paeX* CE12 | *paeX* |  |  |
| *ganA-ganB*<br>GH53 GH42 | *ganA-ganB* |  |  |
| *pelW* PL2 | *pelW* (except C80) |  |  |
| *xynA* GH30 | *xynA* (except C52) |  |  |
| *pehV-pehW-pehX*<br>GH28 GH28 GH28 | *pehV* (except C52) |  |  |
| *pelB-pelC-pelZ*<br>PL1 PL1 PL1 | *pelB* | *pelB-pelC-pelZ* |  |
| *pelN* PL9 | — | *pelN* |  |
| *rhiE* PL4 | — | *rhiE* |  |
| *pemB* CE8 | — | *pemB* |  |
| *pnlG* PL1 | — | *pnlG* |  |
| *pelL-celZ*<br>PL1 GH5 | — |  | *pelL-celZ* |
| *prtA-prtC-prtB-prtF-prtE-prtD-inh-prtG* | — |  |  |

The seven *P. aquatilis* genomes contain a cluster encoding a type II secretion system which is responsible for the specific secretion of extracellular pectinases and of the cellulase CelZ in the genus *Dickeya* [52]. In *P. aquatilis*, it could at least allow the secretion of the pectinases PelB, PelD, PemA, and PaeY (Table 5). In contrast to *Dickeya*, the *P. aquatilis* genomes possess neither genes encoding proteases nor the type I secretion system responsible for their secretion. This is consistent with the absence of protease secretion observed in these strains (Table 1). The genomes of most *P. aquatilis* strains, except LS101 and C111, encode a complete type III secretion system including genes encoding the effector HrpW homologous to PL3 enzymes and the effector DspE belonging to the AvrE family (Table S2).

The number of *luxI-luxR* couples encoding quorum sensing systems dependent on signals of the acyl-homoserine lactone family is variable (Table S2, Table S3). Most of the *P. aquatilis* genomes possess the *smaI-smaR* couple described in strain ATCC39006 [46]. However, *smaI* is a pseudogene in C111 and *smaR* is a pseudogene in C52. Notably, CE90 has a second *luxI-luxR* couple related to the *expI-expR* couple of *Dickeya* (Table S3). In contrast, all the *P. aquatilis* genomes encode the Vfm quorum sensing system previously identified in the genus *Dickeya* which is dependent of the synthesis of non-ribosomal peptides [53]. While the *Dickeya vfm* cluster consists of 26 contiguous genes, the *P. aquatilis* Vfm system consists of 29 genes encoded by two distinct loci (Figure 5, Table S2, Table S3). The major *vfm* locus of *P. aquatilis* contains 26 genes but, in comparison to the *Dickeya* cluster, it shows a different organization of the transcriptional units. Moreover, it lacks the genes *vfmS* and *vfmQ* but contains four additional genes (Figure 5). The second *vfm* locus of *P. aquatilis* contains the genes *vfmJ*, a *vfmF* homolog and *vfmE* encoding the major regulator of the Vfm system. This second locus is adjacent to the *gvp* cluster involved in the production of gas vehicles (Figure 5).

**Fig. 5.**
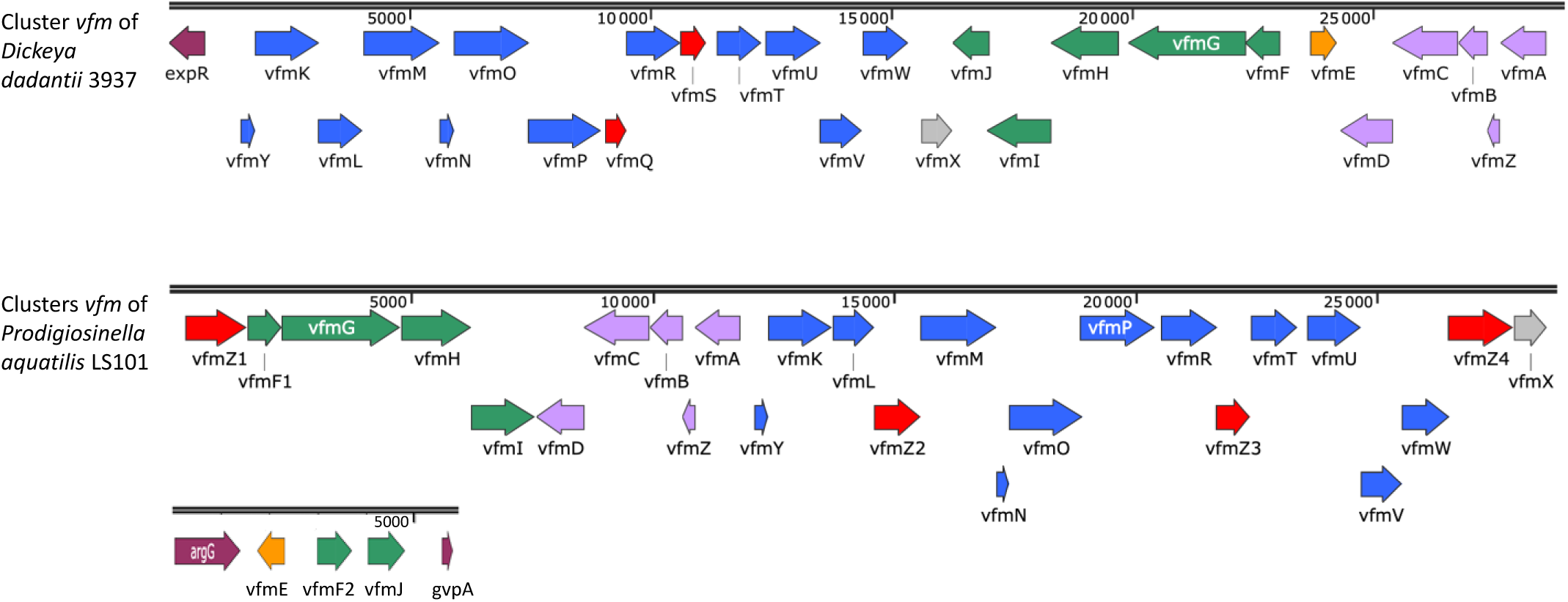
Comparison of the *Dickeya* and *Prodigiosinella vfm* clusters. The *Dickeya vfm* cluster consists of 26 contiguous genes. The *Prodigisinella vfm* genes are located in two separate clusters of 26 and 3 genes, respectively, with a different organization of the transcriptional units which are shown by different colours: blue, *vfmY-vfmW*; green, *vfmF-vfmJ*; purple, *vfmAZBCD*; orange, *vfmE*; grey, *vfmX*. The genes present only in one genus are shown in red. SNAPGENE was used to extract the *vfm* clusters of the 3937 and LS101 genomes.

In conclusion, the isolation of sixteen pectinolytic isolates from Camargue lakes supports the creation of a new genus and the reassignment of the misclassified strain ATCC 39006, as previously suggested by a genome-scale analysis [19]. Phenotypic, phylogenetic and genomic arguments justify the creation of the genus *Prodigiosinella* gen. nov., with *Prodigiosinella aquatilis* as the type species. LS101^T^ (CFBP 8826^T^, LMG 32072^T^) is proposed as the type strain for the new genus and the new species. Moreover, characterization of the different strains led to the identification of two subspecies for which we propose the names *Prodigiosinella aquatilis* subsp *aquatilis* ssp. nov. and *Prodigiosinella aquatilis* subsp *natabilis* ssp. nov., with LS101^T^ (CFBP 8826 ^T^, LMG 32072^T^) and CE70^T^ (CFBP 9054^T^, LMG 32867^T^) as the type strains, respectively. The proposal of these two subspecies is supported by clear genomic and phenotypic differences.

## DESCRIPTION OF THE NEW GENUS, SPECIES AND SUBSPECIES

### Description of *Prodigiosinella* gen. nov

*Prodigiosinella* (Pro.di.gi.o.si.nel’la, N.L. neut. n. prodigiosinum, prodigiosin, the red pigment produced by these bacteria; L. suff. -ella, diminutive ending; N.L. fem. dim. n. *Prodigiosinella* producer of the red pigment prodigiosin).

*Prodigiosinella* members are gram-negative, motile, non-spore forming, facultatively anaerobic bacteria. Cells have average dimensions of 0.6 by 1.7 μm. They are motile and show flotation ability in liquid medium. After 48 h at 30°C on LB medium (5 g.l^-1^ tryptone, 3 g.l^-1^ yeast extract, 5 g.l^-1^ NaCl and 15 g.l^-1^ agar), they form milky-white or pink-colored colonies of 0.5-2 mm in diameter with opaque appearance. They can grow up to 36°C but not at 39°C. They are able to grow in the presence of N-acetyl-D-glucosamine, D-arabinose, L-arabinose, arbutin, D-cellobiose, D-fructose, D-galactose, D-galacturonic acid, gentiobiose, D-gluconic acid, D-glucosamine, α-D-glucose, glycerol, m-inositol, D-mannitol, D-mannose, mucic acid, β-methyl-D-glucoside, pectin, D-ribose, L-rhamnose, D-saccharic acid, salicin, D-sorbitol, sucrose, D-trehalose, D-xylose, γ-amino butyric acid, citric acid, fumaric acid, D-malic acid, L-malic acid, malonic acid, pyruvic acid, and succinic acid. They secrete extracellular pectinases and they produce the red pigment prodigiosin and antibiotic compounds.

Members of this genus include strains LS101^T^ (CFBP 8826^T^, LMG 32072^T^), LS102, LS111, LS112, C52 (CFBP 9053), CE70^T^ (CFBP 9054^T^, LMG 32867^T^), C73, C76, C77, C80 (CFBP 9055), C81, C83, C84, CE90 (CFBP 9051), C110, and C111 (CFBP 9052), isolated from lake water in Camargue and strain ATCC 39006 identified on the basis of its genome sequence. Strains MK01, MK09, and K61 are also *Prodigiosinella aquatilis* members on the basis of their 16S rRNA gene sequence. Strains ATCC 39006 and K61 were isolated in USA (New Jersey and Hawaii, respectively); strains MK01 and MK09 were isolated in Korea.

### Description of Prodigiosinella aquatilis sp. nov

*Prodigiosinella aquatilis* (a.qua’ti.lis. L. fem. adj. *aquatilis* living in water). General description as for the genus.

*P. aquatilis* is able to grow in the presence of N-acetyl-D-glucosamine, D-arabinose, L-arabinose, arbutin, D-cellobiose, D-fructose, D-galactose, D-galacturonic acid, gentiobiose, D-gluconic acid, D-glucosamine, α-D-glucose, glycerol, m-inositol, D-mannitol, D-mannose, mucic acid, β-methyl-D-glucoside, pectin, D-ribose, L-rhamnose, D-saccharic acid, salicin, D-sorbitol, sucrose, D-trehalose, D-xylose, γ-amino butyric acid, citric acid, fumaric acid, D-malic acid, L-malic acid, malonic acid, pyruvic acid, and succinic acid.

The G+C content of the type strain LS101^T^ (CFBP 8826^T^, LMG 32072^T^) genome is 49.1% based on the genome sequence (sequence accession CP128857 (chromosome) and CP128858 (pPrAq1 plasmid).

### Description of Prodigiosinella aquatilis subsp aquatilis ssp. nov

*Prodigiosinella aquatilis* subsp *aquatilis* (a.qua’ti.lis. L. fem. adj. *aquatilis*, living in water).. General description as for the species. *Prodigiosinella aquatilis* subsp *aquatilis* is able to assimilate m-hydroxyphenyl acetic acid, p-hydroxy phenyl acetic acid, tyramine and D.L-octopamine but unable to assimilate D-melibiose and D-raffinose. The type strain is LS101^T^ (CFBP 8826^T^, LMG 32072^T^) and other members of this subspecies are strains LS102, LS111, LS112, C110, C111 (CFBP 9052), and CE90 (CFBP 9051) isolated from lake water in Camargue.

### Description of Prodigiosinella aquatilis subsp natabilis ssp. nov

*Prodigiosinella aquatilis* subsp *natabilis* (na.ˈta.bi.lis. L. fem adj *natabilis*,that can swim or float]. General description as for the species. *Prodigiosinella aquatilis* subsp *natabilis* is able to assimilate D-melibiose, D-raffinose, but unable to assimilate hydroxyphenyl acetic acids, tyramine and octopamine.

CE70^T^ (CFBP 9054^T^, LMG 32867^T^) is the type strain. Other representative isolates of this subspecies include strain ATCC 39006, isolated in channel water in New Jersey (USA), and strains C52 (CFBP 9053), C73, C76, C77, C80 (CFBP 9055), C81, C83 and C84 isolated from lake water in Camargue, France.

## Abbreviations

ANI: average nucleotide identity
BV: Bootstrap value
CAZY: Carbohydrate-Active enZYmes
CE: carbohydrate esterase
CVP medium: crystal violet pectate medium
dDDH: digital DNA–DNA hybridization
GH: glycoside hydrolase
JV: Jackniffe value
LB: Luria–Bertani medium
ML tree: maximum likelihood tree
NCBI: National Center for Biotechnology Information
PGAP: prokaryotic genome annotation pipeline
PL: polysaccharide lyase
T2SS: type II secretion system

## Acknowledgements

We thank Anthony Olivier for its warm welcome in the site of the Foundation Tour du Valat (https://tourduvalat.org/), our lab members for their interest for this work, and Perrine Portier and Claudine Vereecke for strain registration in the CIRM-CFBP and BCCM-LMG collections, respectively.

## REFERENCES

1. Adeolu M, Alnajar S, Naushad S, Gupta RS. Genome-based phylogeny and taxonomy of the *’Enterobacteriales’*: proposal for *Enterobacterales* ord. nov. divided into the families *Enterobacteriaceae*, *Erwiniaceae* fam. nov., *Pectobacteriaceae* fam. nov., *Yersiniaceae* fam. nov., *Hafniaceae* fam. nov., Morganellaceae fam. nov., and Budviciaceae fam. nov. Int J Syst Evol Microbiol 2016; 66: 5575–5599.

2. Kampfer P, Glaeser SP, Nilsson LK, Eberhard T, Hakansson S, et al. Proposal of *Thorsellia kenyensi*s sp. nov. and *Thorsellia kandunguensis* sp. nov., isolated from larvae of Anopheles arabiensis, as members of the family Thorselliaceae fam. nov. Int J Syst Evol Microbiol 2015; 65:444–451.

3. Li M, Liu K, Liu Y, Gao C, Yi X. *Bruguierivorax albus* gen. nov. sp. nov. isolated from mangrove sediment and proposal of *Bruguierivoracaceae* fam. nov. Curr Microbiol 2021; 78:856–866.

4. Lee SD, Kim IS, Choe H, Kim J-S. *Acerihabitans arboris* gen. nov., sp. nov., a new member of the family Pectobacteriaceae isolated from sap drawn from Acer pictum. Int J Syst Evol Microbiol 2021; 71:004674.

5. Bian D-R, Xue H, Wang G-M, Piao C-G, Li Y. *Affinibrenneria salicis* gen. nov. sp. nov. isolated from *Salix matsudana* bark canker. Arch Microbiol 2021; 203: 3473–3481.

6. Brady CL, Cleenwerck I, Denman S, Venter SN, Rodriguez-Palenzuela P, et al. Proposal to reclassify *Brenneria quercina* (Hildebrand and Schroth 1967) Hauben et al. 1999 into a new genus, *Lonsdalea* gen. nov., as *Lonsdalea quercina* comb. nov., descriptions of *Lonsdalea quercina* subsp. *quercina* comb. nov., *Lonsdalea quercina* subsp. *iberica* subsp. nov. and *Lonsdalea quercina* subsp. *britannica* subsp. nov., emendation of the description of the genus *Brenneria*, reclassification of *Dickeya dieffenbachiae* as *Dickeya dadantii* subsp. *dieffenbachiae* comb. nov., and emendation of the description of *Dickeya dadantii*. Int J Syst Evol Microbiol 2012; 62:1592-1602.

7. Samson R, Legendre JB, Christen R, Fischer-Le Saux M, Achouak W, Gardan L. Transfer of *Pectobacterium chrysanthemi* (Burkholder et al. 1953) Brenner et al. 1973 and *Brenneria paradisiaca* to the genus *Dickeya* gen. nov. as *Dickeya chrysanthemi* comb. nov. and *Dickeya paradisiaca* comb. nov. and delineation of four novel species, *Dickeya dadantii* sp. nov., *Dickeya dianthicola* sp. nov., *Dickeya dieffenbachiae* sp. nov. and *Dickeya zeae* sp. nov. Int J Syst Evol Microbiol 2005; 55:1415-1427.

8. Hugouvieux-Cotte-Pattat N, Jacot-des-Combes C, Briolay J, Pritchard L. Proposal for the creation of a new genus *Musicola* gen. nov., reclassification of Dickeya paradisiaca (Samson et al. 2005) as *Musicola paradisiaca* comb. nov. and description of a new species *Musicola keenii* sp. nov. Int J Syst Evol Microbiol 2021; 71(10):005037.

9. Hauben L, Moore ERB, Vauterin L, Steenackers M, Mergaert J, et al. Phylogenetic position of phytopathogens within the *Enterobacteriaceae*. Syst Appl Microbiol 1998; 21:384–397.

10. Sutra L, Christen R, Bollet C, Simoneau P, Gardan L. S*amsonia erythrinae* gen. nov., sp. nov., isolated from bark necrotic lesions of Erythrina sp., and discrimination of plant pathogenic Enterobacteriaceae by phenotypic features. Int J Syst Evol Microbiol (2001; 51:1291–1304.

11. Nadal-Jimenez P, Siozios S, Halliday N, Cámara M, Hurst GDD. *Symbiopectobacterium purcellii*, gen. nov., sp. nov., isolated from the leafhopper Empoasca decipiens. Int J Syst Evol Microbiol 2022; 72(6):005440

12. Hugouvieux-Cotte-Pattat N, Condemine G, Gueguen E, Shevchik VE. *Dickeya* Plant Pathogens. In eLS. John Wiley & Sons, Hoboken, NJ 2020: 1–10.

13. Van Gijsegem F, Hugouvieux-Cotte-Pattat N, Kraepiel Y, Lojkowska E, Moleleki LN, et al. Molecular interactions of Pectobacterium and Dickeya with plants. In: Van Gijsegem F, van der Wolf J and Toth IK (eds). Plant Diseases Caused by Dickeya and Pectobacterium Species. New York: Springer; 2021. pp. 85–147.

14. Parkinson N, DeVos P, Pirhonen M, Elphinstone J. *Dickeya aquatica* sp. nov., isolated from waterways. Int J Syst Evol Microbiol 2014; 64:2264–2266.

15. Hugouvieux-Cotte-Pattat N, Jacot-des-Combes C, Briolay J.. *Dickeya lacustris* sp. nov., a water-living pectinolytic bacterium isolated from lakes in France. Int J Syst Evol Microbiol 2019; 69: 721–726.

16. Oulghazi S, Pédron J, Cigna J, Lau YY, Moumni M, et al. *Dickeya undicola* sp. nov., a novel species for pectinolytic isolates from surface waters in Europe and Asia. Int J Syst Evol Microbiol 2019; 69: 2440–2444.

17. Helias V, Hamon P, Huchet E, Wolf JVD, Andrivon D. Two new effective semiselective crystal violet pectate media for isolation of *Pectobacterium* and *Dickeya*. Plant Pathol 2012; 61: 339–345.

18. Duprey A, Taib N, Leonard S, Garin T, Flandrois JP, et al. The phytopathogenic nature of *Dickeya aquatica* 174/2 and the dynamic early evolution of *Dickeya* pathogenicity. Environ Microbiol 2019; 21: 2809–2835.

19. Cigna J, Dewaegeneire P, Beury A, Gobert V, Faure D. A *gapA* PCR sequencing assay for identifying the *Dickeya* and *Pectobacterium* potato pathogens. Plant Disease 2017; 101: 1278–1282.

20. Gouy M, Guindon S. Gascuel O. SeaView version 4: a multiplatform graphical user interface for sequence alignment and phylogenetic tree building. Mol Biol Evol 2010; 27:221–224.

21. Edgar RC. MUSCLE: multiple sequence alignment with high accuracy and high throughput. Nucleic Acids Res 2004; 32:1792–1797.

22. Minh BQ, Schmidt HA, Chernomor O, Schrempf D, Woodhams MD, et al. IQ-TREE 2: New models and efficient methods for phylogenetic inference in the genomic era. Mol Biol Evol 2020; 37:1530–1534.

23. Kalyaanamoorthy BQ, Minh Wong TKF, von Haeseler A, Jermiin LS. ModelFinder: Fast model selection for accurate phylogenetic estimates. Nat Methods 2017; 14:587–589.

24. Lane DJ, Pace B, Olsen GJ, Stahl DA, Sogin ML et al. Rapid determination of 16S ribosomal RNA sequences for phylogenetic analyses. Proc Natl Acad Sci USA 1985; 82: 6955–6959.

25. Parker WL, Rathnum ML, Wells JS, Trejo WH, Principe PA, Sykes RB. SQ 27,860, a simple carbapenem produced by species of *Serratia* and *Erwinia*. J Antibiot (Tokyo) 1982; 35: 653–660.

26. Fineran PC, Iglesias Cans MC, Ramsay JP, Wilf NM, Cossyleon D et al. Draft genome sequence of *Serratia* sp. strain ATCC 39006, a model bacterium for analysis of the biosynthesis and regulation of prodigiosin, a carbapenem, and gas vesicles. Genome Announc 2013; 1(6):e01039–13.

27. Katoh S. MAFFT multiple sequence alignment software version 7: improvements in performance and usability. Mol Biol Evol 2013; 30:772–780.

28. Criscuolo A, Gribaldo S. BMGE (Block Mapping and Gathering with Entropy): a new software for selection of phylogenetic informative regions from multiple sequence alignments. BMC Evol Biol 2010; 10:210.

29. Coutinho TA, Yoon Shin G, van der Waals J. Dickeya. In Bergey’s Manual of Systematics of Archaea and Bacteria. John Wiley & Sons. 2021; 10.1002/9781118960608.gbm02014

30. Coutinho TA, Yoon Shin G, van der Waals J. Pectobacterium. In Bergey’s Manual of Systematics of Archaea and Bacteria. John Wiley & Sons. 2021; 10.1002/9781118960608.gbm01158

31. Bochner BR. Global phenotypic characterization of bacteria. FEMS Microbiol Rev 2009; 33: 191–205.

32. Tashiro Y, Monson RE, Ramsay JP, Salmond GP. Molecular genetic and physical analysis of gas vesicles in buoyant enterobacteria. Environ Microbiol 2016; 18: 1264–1276.

33. Hugouvieux-Cotte-Pattat N, Jacot-des-Combes C, Briolay J. Genomic characterization of a pectinolytic isolate of *Serratia oryzae* isolated from lake water. J Genomics 2019; 7: 64–72.

34. Kolmogorov M, Yuan J, Lin Y, Pevzner P. Assembly of long error-prone reads using repeat graphs. Nature Biotechnology 2019; 37: 540–546.

35. Wick RR, Schultz MB, Zobel J, Holt KE. Bandage: interactive visualisation of de novo genome assemblies. Bioinformatics 2015; 31: 3350–3352.

36. Langmead B, Trapnell C, Pop M, Salzberg SL. Ultrafast and memory-efficient alignment of short DNA sequences to the human genome. Genome Biol 2009; 10:R25.

37. Li H, Handsaker B, Wysoker A, Fennell T, Ruan J, Homer N, Marth G, Abecasis G, Durbin R, and 1000 Genome Project Data Processing Subgroup, The Sequence alignment/map (SAM) format and SAMtools, Bioinformatics 2009; 25: 2078–2079.

38. Walker BJ, Abeel T, Shea T, Priest M, Abouelliel A, Sakthikumar S, Cuomo CA, Zeng Q, Wortman J, Young SK, Earl AM. Pilon: An integrated tool for comprehensive microbial variant detection and genome assembly improvement. PLoS ONE 2014; 9(11): e112963.

39. Goris J, Konstantinidis KT, Klappenbach JA, Coenye T, Vandamme P, et al. DNA-DNA hybridization values and their relationship to whole-genome sequence similarities. Int J Syst Evol Microbiol 2007; 57: 81–91.

40. Meier-Kolthoff JP, Auch AF, Klenk HP, Göker M. Genome sequence-based species delimitation with confidence intervals and improved distance functions. BMC Bioinformatics 2013; 14: 60.

41. Meier-Kolthoff JP, Hahnke RL, Petersen J, Scheuner C, Michael V et al. Complete genome sequence of DSM 30083(T), the type strain (U5/41(T)) of *Escherichia coli*, and a proposal for delineating subspecies in microbial taxonomy. Stand Genomic Sci 2014; 9: 2.

42. Li W, O’Neill KR, Haft DH, DiCuccio M, Chetvernin V, et al. Expanding the Prokaryotic Genome Annotation Pipeline reach with protein family model curation. Nucleic Acids Res 2021; 49(D1):D1020–D1028.

43. Jauffrit F, Penel S, Delmotte S, Rey C, de Vienne DM, et al. RiboDB Database: A Comprehensive Resource for Prokaryotic Systematics. Mol Biol Evol 2016; 33: 2170–2172.

44. Harris AKP, Williamson NR, Slater H, Cox A, Abbasi S, et al. The *Serratia* gene cluster encoding biosynthesis of the red antibiotic, prodigiosin, shows species- and strain-dependent genome context variation. Microbiology 2004; 150: 3547–3560.

45. McGowan SJ, Holden MT, Bycroft BW, Salmond GP. Molecular genetics of carbapenem antibiotic biosynthesis. Antonie Van Leeuwenhoek 1999; 75: 135–141.

46. Ramsay JP, Williamson NR, Spring DR, Salmond GP. A quorum-sensing molecule acts as a morphogen controlling gas vesicle organelle biogenesis and adaptive flotation in an enterobacterium. Proc Natl Acad Sci USA. 2011; 108: 14932–14937.

47. Williams DJ, Grimont PAD, Cazares A, Grimont F, Ageron E, et al. The genus *Serratia* revisited by genomics. Nat Commun. 2022; 13:5195.

48. Diaz E, Ferrandez A, Prieto MA, Garcia JL. Biodegradation of aromatic compounds by *Escherichia coli*. Microbiol Mol Biol Rev. 2001; 65: 523–569.

49. Luengo JM, Olivera ER. Catabolism of biogenic amines in *Pseudomonas* species. Environ Microbiol 2020; 22: 1174–1192.

50. Hugouvieux-Cotte-Pattat N, Condemine G, Shevchik VE. Bacterial pectate lyases, structural and functional diversity. Environ Microbiol Rep 2014; 6: 427–440.

51. Drula E, Garron ML, Dogan S, Lombard V, Henrissat B, Terrapon N. The carbohydrate-active enzyme database: functions and literature. Nucleic Acids Research 2022; 50: D571–D577.

52. Pineau C, Guschinskaya N, Robert X, Gouet P, Ballut L, Shevchik VE. Substrate recognition by the bacterial type II secretion system: more than a simple interaction. Mol Microbiol 2014; 94: 126–140.

53. Hugouvieux-Cotte-Pattat N, Royer M, Gueguen E, Le Guen P, Süssmuth RD, Reverchon S, Cociancich S. Specificity and genetic polymorphism in the Vfm quorum sensing system of plant pathogenic bacteria of the genus *Dickeya*. Environ Microbiol. 2022; 24: 1467–1483.

