## supplementary data for "Description of a new genus of the *Pectobacteriaceae* family isolated from lake water in France; *Prodigiosinella aquatilis* gen. nov. sp. nov. includes two subspecies *Prodigiosinella aquatilis* subsp. *aquatilis* ssp. nov. and *Prodigiosinella aquatilis* subsp. *natabilis* ssp. nov"

### Fig. S1. Growth of six *Prodigiosinella* strains at different temperatures

To analyse the growth temperature, bacterial cultures were performed in LB medium in the range of 21 to 45°C. The cell density was estimated by measuring the optical density at 600 nm ($OD_{600}$) after 24 h. *D. dadantii* 3937 and *D. lacustris* S29 were used for comparison.

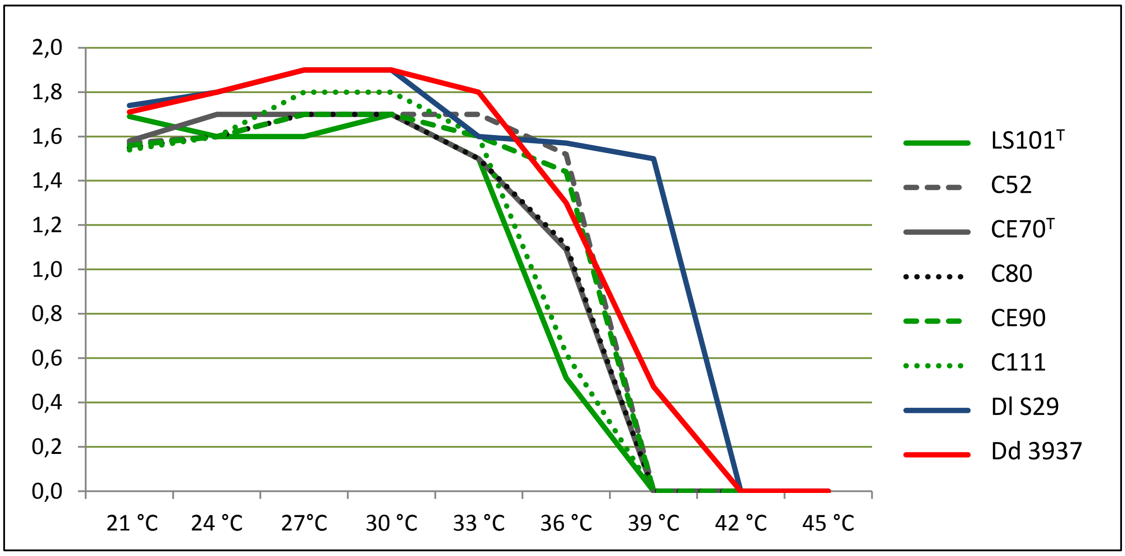

*Prodigiosinella aquatilis* subsp. *aquatilis* (group G1) LS101$^{T}$, CE90, C111 (green lines)
*Prodigiosinella aquatilis* subsp. *natabilis* (group G2) CE70$^{T}$, C52, C80 (grey/black lines)
*Dickeya lacustris* S29$^{T}$ (blue line)
*Dickeya dadantii* 3937 (red line)

**Fig. S2. Maximum likelihood phylogeny of the *Pectobacteriaceae* based on ribosomal protein genes.** The tree shows the relationships between the six Camargue isolates, strain ATCC 39006, and *Pectobacteriaceae* genera (87 strains, 19,311 nucleotide positions). The tree is rooted with representatives of other *Enterobacterales* families. It was inferred with IQ-TREE (SYM+I+I+R5 evolutionary model). The scale bar represents the average number of substitutions per site. Grey circles correspond to BV = 100% computed by IQ-TREE (1,000 replicates).

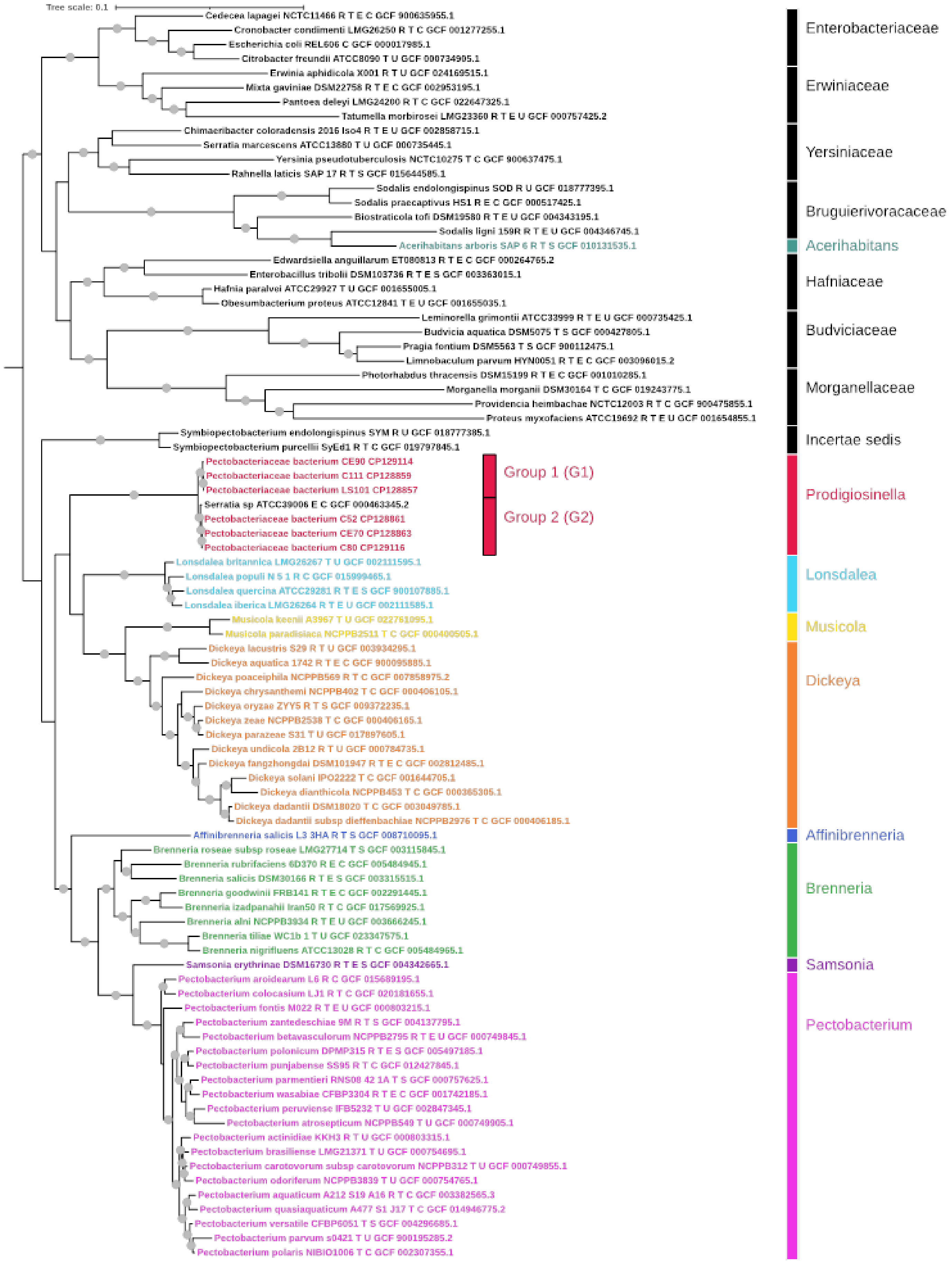

**Table S1. Metabolic capacities of 3 strains from each *Prodigiosinella* subspecies**

These data were obtained using Biolog plates PM1 and PM2A. Bacterial growth was determined after 48h at 30°C, by measurement of optical density (OD) at λ=590nm: -, indicates OD <0.2; w, indicates 0.2≤OD≤0.5; +, indicates OD> 0.5. The characters showing variations among the two subspecies are coloured (green, characters specific of P. *aquatilis aquatilis*; red, characters specific of P. *aquatilis natabilis*.

| Strains | LS101^T A6420 | C111 A6483 | CE90 A6482 | C52 A6477 | C80 A6480 | CE70^T A6479 |
|---|---|---|---|---|---|---|
| ***Prodigiosinella aquatilis* subspecies** | ***aquatilis*** | ***aquatilis*** | ***aquatilis*** | ***natabilis*** | ***natabilis*** | ***natabilis*** |
| **L-Arabinose** | **+** | **+** | **+** | **+** | **w** | **+** |
| **N-Acetyl-D-glucosamine** | **+** | **+** | **+** | **+** | **+** | **+** |
| **D-Saccharic acid (glucaric acid)** | **+** | **+** | **+** | **+** | **+** | **+** |
| **Succinic acid** | **+** | **w** | **+** | **+** | **w** | **+** |
| **D-Galactose** | **+** | **+** | **+** | **+** | **+** | **+** |
| **L-Aspartic acid** | **+** | **+** | **+** | **+** | **+** | **+** |
| L-Proline | - | - | - | w | w | - |
| D-Alanine | - | - | - | w | w | w |
| **D-Trehalose** | **+** | **+** | **+** | **+** | **+** | **+** |
| **D-Mannose** | **+** | **+** | **+** | **+** | **+** | **+** |
| Dulcitol | - | - | - | - | - | - |
| D-Serine | - | - | - | - | - | - |
| **D-Sorbitol** | **+** | **+** | **+** | **+** | **+** | **+** |
| **Glycerol** | **+** | **+** | **+** | **+** | **+** | **+** |
| L-Fucose | - | - | - | - | - | - |
| D-Glucuronic acid | w | - | - | - | - | - |
| **D-Gluconic acid** | **+** | **+** | **+** | **+** | **+** | **+** |
| D,L-α-Glycerol- phosphate | - | - | - | w | w | w |
| **D-Xylose** | **+** | **+** | **+** | **+** | **w** | **+** |
| L-Lactic acid | - | w | + | + | w | + |
| Formic acid | - | - | w | + | w | + |
| **D-Mannitol** | **+** | **+** | **+** | **+** | **+** | **+** |
| L-Glutamic acid | - | - | w | + | w | + |
| D-Glucose-6-phosphate | - | - | - | - | - | - |
| D-Galactonic acid-γ-lactone | - | - | - | - | - | - |
| **D,L-Malic acid** | **+** | **+** | **+** | **+** | **+** | **+** |
| **D-Ribose** | **+** | **+** | **+** | **+** | **w** | **w** |
| Tween 20 | - | - | - | - | - | - |
| **L-Rhamnose** | **+** | **+** | **+** | **+** | **+** | **+** |
| **D-Fructose** | **+** | **+** | **+** | **+** | **+** | **+** |
| Acetic acid | w | - | - | w | w | + |
| **α-D-Glucose** | **+** | **+** | **+** | **+** | **+** | **+** |
| Maltose | w | w | - | w | - | - |
| **D-Melibiose** | **-** | **-** | **-** | **+** | **+** | **+** |
| Thymidine | **-** | **-** | **-** | **w** | **w** | **w** |
| **L-Asparagine** | **+** | **+** | **+** | **+** | **+** | **+** |
| **D-Aspartic acid** | **+** | **+** | **+** | **+** | **w** | **+** |
| D-Glucosaminic acid | w | w | w | w | - | - |
| 1,2-Propanediol | - | - | - | - | - | - |
| Tween 40 | - | - | - | - | - | - |

| | | | | | | |
|---|---|---|---|---|---|---|
| α-Keto-glutaric acid | - | - | - | - | - | - |
| α-Keto-butyric acid | - | - | - | - | - | - |
| α-Methyl-D-galactoside | - | - | - | - | w | w |
| α-D-Lactose | - | - | - | - | - | - |
| Lactulose | - | - | - | - | - | - |
| **Sucrose** | **+** | **+** | **+** | **+** | **+** | **+** |
| Uridine | - | - | - | - | - | - |
| L-Glutamine | - | - | - | w | - | w |
| m-Tartaric Acid | - | - | - | w | - | - |
| D-Glucose-1-Phosphate | + | + | + | - | + | + |
| D-Fructose-6-Phosphate | - | - | - | - | - | - |
| Tween 80 | - | - | - | - | - | - |
| α-Hydroxy glutaric acid-γ-lactone | - | - | - | - | - | - |
| α-Hydroxy butyric acid | - | - | - | - | - | - |
| **β-Methyl-D-glucoside** | **+** | **+** | **+** | **w** | **+** | **+** |
| Adonitol | - | - | - | - | - | - |
| Maltotriose | + | w | - | - | - | - |
| 2-Deoxy adenosine | - | - | - | - | - | - |
| Adenosine | - | - | - | - | - | - |
| Glycyl-L-aspartic acid | - | - | - | - | - | - |
| **Citric acid** | **+** | **+** | **+** | **+** | **+** | **+** |
| **m-Inositol** | **+** | **+** | **+** | **+** | **+** | **+** |
| D-Threonine | - | - | - | - | - | - |
| **Fumaric acid** | **+** | **+** | **+** | **+** | **+** | **+** |
| **Bromo succinic acid** | **+** | **w** | **+** | **w** | **+** | **+** |
| Propionic acid | - | - | - | - | - | - |
| **Mucic acid (galactaric acid)** | **+** | **+** | **+** | **+** | **+** | **+** |
| Glycolic acid | - | - | - | - | - | - |
| Glyoxylic acid | - | - | - | - | - | - |
| **D-Cellobiose** | **+** | **+** | **+** | **+** | **+** | **+** |
| Inosine | - | - | - | - | - | - |
| Glycyl-L-glutamic acid | - | - | - | - | - | - |
| Tricarballylic acid | - | - | - | - | - | - |
| **L-Serine** | **+** | **+** | **w** | **+** | **+** | **+** |
| L-Threonine | - | - | - | - | - | - |
| L-Alanine | - | - | - | w | w | w |
| L-Alanyl-glycine | - | - | - | - | - | - |
| Acetoacetic acid | - | - | - | - | - | - |
| N-Acetyl-β-D-mannosamine | - | - | - | - | - | - |
| Mono methyl succinate | - | - | - | - | - | w |
| **Methyl pyruvate** | **+** | **w** | **w** | **+** | **w** | **+** |
| **D-Malic acid** | **+** | **w** | **w** | **+** | **w** | **+** |
| **L-Malic acid** | **+** | **+** | **+** | **+** | **+** | **+** |
| Glycyl-L-proline | - | - | - | - | - | - |
| **p-Hydroxy phenyl acetic acid** | **+** | **+** | **w** | - | - | - |
| **m-Hydroxy phenyl acetic acid** | **w** | **w** | **w** | - | - | - |
| **Tyramine** | **+** | **w** | **w** | - | - | - |
| D-Psicose | - | - | - | - | - | - |
| L-Lyxose | - | - | - | - | - | - |
| Glucuronamide | - | - | - | - | - | - |
| **Pyruvic acid** | **+** | **+** | **+** | **+** | **+** | **+** |

| | | | | | | |
|---|---|---|---|---|---|---|
| L-Galactonic acid-γ-lactone | + | w | - | + | + | + |
| **D-Galacturonic acid** | **+** | **+** | **+** | **+** | **+** | **+** |
| Phenylethyl-amine | - | - | - | - | - | - |
| 2-Aminoethanol | - | - | - | - | - | - |
| Chondroitin sulfate C | - | - | - | - | - | - |
| α-Cyclodextrin | - | - | - | - | - | - |
| β-Cyclodextrin | - | - | - | - | - | - |
| γ-Cyclodextrin | - | - | - | - | - | - |
| Dextrin | - | - | - | - | - | - |
| Gelatin | - | - | - | - | - | w |
| Glycogen | - | - | - | - | - | - |
| Inulin | - | - | - | - | - | - |
| Laminarin | - | - | - | - | - | - |
| Mannan | - | - | - | - | - | - |
| **Pectin** | **+** | **+** | **+** | **+** | **+** | **+** |
| N-Acetyl-D-galactosamine | - | - | - | - | - | - |
| N-Acetyl-neuraminic acid | - | - | - | - | - | - |
| β-D-Allose | + | + | - | + | w | w |
| Amygdalin | - | - | - | - | - | - |
| D-Arabinose | + | + | - | + | - | - |
| D-Arabitol | - | - | - | - | - | - |
| L-Arabitol | - | - | - | - | - | - |
| **Arbutin** | **+** | **+** | **+** | **+** | **+** | **+** |
| 2-Deoxy-D-ribose | - | - | - | - | - | - |
| I-Erythritol | - | - | - | - | - | - |
| D-Fucose | - | - | - | - | - | - |
| 3-0-β-D-Galacto-pyranosyl-D-arabinose | w | - | - | - | - | - |
| **Gentiobiose** | **+** | **+** | **w** | **+** | **+** | **+** |
| L-Glucose | - | - | - | - | - | - |
| Lactitol | - | - | - | - | - | - |
| D-Melezitose | - | - | - | - | - | - |
| Maltitol | - | - | - | - | - | - |
| a-Methyl-D-glucoside | - | - | - | - | - | - |
| β-Methyl-D-galactoside | - | - | - | - | - | - |
| 3-Methyl Glucose | - | - | - | - | - | - |
| β-Methyl-D-glucuronic acid | - | - | - | - | - | - |
| α-Methyl-D-mannoside | - | - | - | - | - | - |
| β-Methyl-D-xyloside | - | - | - | - | - | - |
| Palatinose | - | - | - | - | - | - |
| **D-Raffinose** | **-** | **-** | **-** | **+** | **+** | **+** |
| **Salicin** | **+** | **+** | **+** | **+** | **+** | **+** |
| Sedoheptulosan | - | - | - | - | - | - |
| L-Sorbose | - | - | - | - | - | - |
| Stachyose | - | - | - | - | - | - |
| D-Tagatose | - | - | - | - | - | - |
| Turanose | - | - | - | - | - | - |
| Xylitol | - | - | - | - | - | - |
| N-Acetyl-D-glucosaminitol | - | - | - | - | - | - |
| **γ-Amino butyric acid** | **+** | **+** | **+** | **+** | **w** | **+** |
| δ-Amino valeric acid | - | - | - | - | - | - |
| Butyric acid | - | - | - | - | - | - |

| | | | | | | |
|---|---|---|---|---|---|---|
| Capric acid | - | - | - | - | - | - |
| Caproic acid | - | - | - | - | - | - |
| Citraconic acid | - | - | - | - | - | - |
| Citramalic acid | - | - | - | - | - | - |
| **D-Glucosamine** | **+** | **+** | **+** | **+** | **+** | **+** |
| 2-Hydroxy benzoic acid | - | - | - | - | - | - |
| 4-Hydroxy benzoic acid | - | - | - | - | - | - |
| β-Hydroxy butyric acid | - | - | - | - | - | - |
| γ-Hydroxy butyric acid | - | - | - | - | - | - |
| a-Keto-valeric acid | - | - | - | - | - | - |
| Itaconic acid | - | - | - | - | - | - |
| 5-Keto-D-gluconic acid | w | - | - | w | - | - |
| D-Lactic acid methyl ester | w | w | - | w | - | - |
| **Malonic acid** | **w** | **w** | **w** | **w** | **w** | **+** |
| Melibionic acid | - | - | - | - | - | - |
| Oxalic acid | - | - | - | - | - | - |
| Oxalomalic acid | - | - | - | - | - | - |
| Quinic acid | - | - | - | - | - | - |
| D-Ribono-1,4-lactone | - | - | - | - | - | - |
| Sebacic acid | - | - | - | - | - | - |
| Sorbic acid | - | - | - | - | - | - |
| Succinamic acid | - | w | + | w | w | + |
| D-Tartaric acid | - | - | - | - | - | - |
| L-Tartaric acid | - | - | - | - | - | - |
| Acetamide | - | - | - | - | - | - |
| L-Alaninamide | - | - | - | - | - | - |
| N-Acetyl-L-glutamic acid | - | - | - | - | - | - |
| L-Arginine | - | - | - | - | - | - |
| Glycine | - | - | - | - | - | - |
| L-Histidine | - | - | - | - | - | - |
| L-Homoserine | - | - | - | - | - | - |
| Hydroxy-L-proline | - | - | - | - | - | - |
| L-Isoleucine | - | - | - | - | - | - |
| L-Leucine | - | - | - | - | - | - |
| L-Lysine | - | - | - | - | - | - |
| L-Methionine | - | - | - | - | - | - |
| L-Ornithine | - | - | - | - | - | - |
| L-Phenylalanine | - | - | - | - | - | - |
| L-Pyroglutamic acid | - | - | - | - | - | - |
| L-Valine | - | - | - | - | - | - |
| D,L-Carnitine | - | - | - | - | - | - |
| Sec-Butylamine | - | - | - | - | - | - |
| **D.L-Octopamine** | **+** | **+** | **w** | **-** | **-** | **-** |
| Putrescine | - | - | - | - | - | - |
| Dihydroxy Acetone | - | - | - | - | - | - |
| 2,3-Butanediol | - | - | - | - | - | - |
| 2,3-Butanedione | - | - | - | - | - | - |
| 3-Hydroxy 2-Butanone | - | - | - | - | - | - |

**Table S2. Gene distribution within the *Prodigiosinella aquatilis* genomes.** (Excel table)

The table shows the presence/absence of genes encoding specific functions of *Prodigiosinella* strains, such as the biosynthesis of prodigiosin and carbapenem, the production of gas vehicles, and the assimilation of hydroxyphenyl acetic acids. It also gives the presence/absence of genes encoding plant cell wall degrading enzymes, type II and type III secretions systems and quorum sensing systems. Genes were considered as present if identity of the encoded proteins was higher than 60% of full-length amino acids sequence.

**Table S3. Gene clusters involved in quorum sensing regulation in *Prodigiosinella aquatilis* strains.**

These regulations imply couples of the LuxI-LuxR family and the Vfm system previously described in the genus *Dickeya*. Most *P. aquatilis* strains possess the *smaI-smaR* couple described in ATCC39006; CE90 has a second *luxI-luxR* couple related to the *expI-expR* couple of *Dickeya.*
The *Dickeya vfm* cluster consists of 26 contiguous genes. The *Prodigisinella vfm* genes are located in two separate clusters of 26 and 3 genes, respectively, with a different genetic organization. Genes of the *vfm* cluster shown in red are present only in *only one* species (*vfmQ* and *vfmS* in *D. dadantii*, *vfmZ1, vfmZ2, vfmZ3* and *vfmZ4* in *P. aquatilis*).

| Strains | LuxI-LuxR family | Vfm system |
|---|---|---|
| *Dickeya dadantii* 3937 | *expI-expR* | *vfmAZBCDEFGHIJXYKLMNOP Q R S TW* |
| *P. aquatilis* subsp. *aquatilis* LS101$^T$ | *smaI-smaR* | *vfmAZBCD Z*$_1$ *F*$_1$ *GHIJXYKL Z*$_2$ *MNOPR Z*$_3$ *TW Z*$_4$<br>*vfmEF*$_2$ *J* |
| *P. aquatilis* subsp. *natabilis* CE70$^T$, C80, ATCC 39006 | | |
| *P. aquatilis* subsp. *aquatilis* CE90 | *smaI-smaR, expI-expR* | |
| *P. aquatilis* subsp. *natabilis* C52 | *smaI* | |
| *P. aquatilis* subsp. *aquatilis* C111 | – | |
